# Structural insights into antibody responses against influenza A virus in its natural reservoir

**DOI:** 10.64898/2026.03.11.711171

**Authors:** Huibin Lv, Walter N. Harrington, Wenkan Liu, Dalia Naser, Yang Wei Huan, Emma Thames, Pradeep Chopra, Tossapol Pholcharee, Patrick Seiler, Edgardo Ayala, Alyssa Monterroso, Wei Ji, Qi Wen Teo, Akshita B. Gopal, Emily X. Ma, Douglas C. Wu, Madison R. Ardagh, Arjun Mehta, Jessica J. Huang, Meixuan Tong, Garrett Honzay, Brendan Shirkey, Jenna J. Guthmiller, Geert-Jan Boons, Beth M. Stadtmueller, Richard J. Webby, Nicholas C. Wu

## Abstract

While influenza A virus undergoes rapid antigenic drift in humans, at least some subtypes, such as H3, have relatively stable antigenicity in natural waterfowl reservoirs, despite the presence of immune pressure. However, the underlying mechanisms remain poorly understood. This study identified and characterized 187 antibodies to H3 hemagglutinin from experimentally infected mallard ducks, 18 of which were further analyzed by cryo-EM. Compared with human H3 antibodies, duck H3 antibodies exhibited higher glycan-binding propensity, more balanced immunodominance hierarchy, and targeted distinct epitopes. Other unique features of duck H3 antibodies included a convergent CDR H3-independent heavy chain-only binding mode and an N-glycosylated CDR H3 as decoy receptor. By annotating duck immunoglobulin germline genes, we also demonstrated the importance of gene conversion in duck H3 antibodies. Overall, our findings provide insights into how millennia of coevolution have shaped the interplay between influenza A virus antigenic drift and antibody responses in the natural reservoir.

## INTRODUCTION

Despite being discovered almost a century ago^1,2^, influenza A virus remains a global public health concern. The major antigen of influenza A virus, hemagglutinin (HA), consists of 19 known subtypes (H1-H19)^3^. While current seasonal influenza epidemics are caused by H1 and H3 subtypes, other zoonotic subtypes occasionally spill over into humans and continually pose pandemic threats^4,5^. As the natural reservoir of influenza A virus, Anseriformes (ducks, geese, and swans) and Charadriiformes (shorebirds), are commonly exposed to different, mostly low-pathogenic, subtypes of influenza A virus in the intestinal tract^6,7^. Notably, dabbling ducks (feeding primarily in shallow water), such as mallards, are infected with influenza A virus more frequently than other waterfowl species, including diving ducks (feeding in deep water)^7^. Therefore, dabbling ducks serve as a primary reservoir host for influenza A virus. It is estimated that the common ancestor of avian influenza A virus subtypes arose thousands of years ago^8,9^. This long evolutionary history has likely shaped the immune response of dabbling ducks, as demonstrated by their minimal clinical signs observed during most influenza A virus infections^10–13^. The limited pathogenesis of influenza A virus in dabbling ducks has been attributed to their robust innate immune responses^14–16^, especially against highly pathogenic strains^17^. This notion is further supported by a genomic analysis showing expansions of their β-defensin and butyrophilin-like repertoires^18^. In contrast, it is largely unclear how thousands of years of coevolution have molded the antibody responses of dabbling ducks against influenza A virus.

The HA of human influenza A virus undergoes rapid antigenic drift due to the positive selection pressure imposed by antibody responses. Antigenic drift of human H3 HA is mainly caused by mutations in the five major antigenic sites (A-E) in the head domain^19–21^, which are located above the immunosubdominant HA stem domain^3^. Some of these antigenic mutations also lead to the introduction of N-glycosylation sites^21,22^, resulting in an increasing number of N-glycans in human H3 HA over time^23,24^. On the other hand, duck H3 HA is rather antigenically stable, despite its sequence diversity^25–28^. For example, duck H3 HA exhibits highly similar antigenicity across more than 20 years, as determined by the binding profiles of a panel of monoclonal antibodies^27^. Consistently, a comparative analysis of influenza evolution in different host species reveals an exceptionally low ratio of non-synonymous to synonymous substitutions (dN/dS) in avian H1 and H3 HAs^29^. At the same time, dabbling ducks can mount neutralizing antibody responses against HA following influenza A virus infection^10,30–33^, both in the blood and intestinal tract^10,34^. Moreover, dabbling ducks have high seroprevalence^35–37^ and impose immune pressure on circulating influenza A virus through homosubtypic and heterosubtypic immunity^38^. Although ecological and epidemiological factors could contribute to the disparity in antigenic evolution of influenza A virus between ducks and humans^27,39^, the distinct molecular characteristics of their antibody responses likely also play a role.

Chickens have historically served as the conventional model for studying avian antibody responses, especially since B cells were first discovered in chickens^40^. Chicken antibodies utilize a single functional immunoglobulin heavy variable (*IGHV*) gene and a single functional immunoglobulin lambda variable (*IGLV*) gene, which are diversified through gene conversion from pseudogenes^41,42^. However, this single functional gene configuration is not universal across all avian species. For example, multiple functional *IGLV* genes are present in dabbling ducks, including the Muscovy duck^43^ and the Pekin duck^44^. Substantial differences in the immunoglobulin loci between ducks and chickens are not unexpected, given their approximately 90 million years of divergence^45,46^, an evolutionary distance similar to that between humans and rodents^47,48^. Nevertheless, while the immunoglobulin germline genes of chickens have been systematically documented^49^, those of ducks have not. Additionally, to the best of our knowledge, only two heavy chain sequences of duck monoclonal antibodies have been reported to date, and their specificities remain uncharacterized^50^. Consequently, there is a lack of molecular understanding of duck antibody responses, including those against the influenza A virus.

In this study, we performed an in-depth molecular analysis of mallard duck antibody responses against influenza A virus infection. By leveraging whole-genome sequencing, single-cell sequencing, and phage display antibody screening, we not only annotated the mallard duck immunoglobulin germline genes, but also identified and characterized 187 antibodies to H3 HA from experimentally infected mallard ducks. Furthermore, we determined cryogenic electron microscopy (cryo-EM) structures for 18 mallard duck antibodies in complex with HA. Our results revealed an unexpected HA glycan-binding propensity among mallard duck antibodies, indicating that avian influenza A virus may not be able to escape mallard duck antibody responses by accumulating N-glycosylation sites. At the same time, the mallard duck antibody response to HA head domain displayed a relatively balanced immunodominance hierarchy, which could increase the genetic barrier for antibody escape. These observations helped explain the antigenic stability of influenza A virus H3 subtype in its natural reservoir despite the presence of antibody responses^25–28,35–37,51^. Moreover, we discovered a heavy chain-only but complementarity-determining region (CDR) H3-independent binding mode in mallard duck antibody responses, as well as a mallard duck antibody that can partially function as a decoy receptor via an N-glycosylated CDR H3. These findings uncover the differences between duck and human antibody responses and their potential to differentially impact influenza evolution.

## RESULTS

### Annotation of duck immunoglobulin germline genes

We first aimed to analyze the cell-type composition of mallard duck peripheral blood. Single-cell sequencing analysis of the peripheral blood mononuclear cells (PBMCs) from a wild mallard duck (*Anas platyrhynchos*) showed that 2.4% of PBMCs were B cells (**Figure 1A-B**). We further performed whole-genome sequencing of a mallard duck using PacBio long-read sequencing. The resulting genome assembly achieved an N50 of 20 Mb with 97% genome completeness and around 98x read coverage (**Figure S1A-B**), allowing reconstruction of the immunoglobulin heavy and light chain loci. Subsequently, we identified immunoglobulin germline genes through BLAST searches and locating the recombination signal sequences (RSS) (**Figure 1C-D and Table S1**). Within the heavy chain locus, we identified a single functional immunoglobulin heavy variable (*IGHV*) gene (*IGHV1-1*) and 57 pseudogenes (*IGHV1-2* to *IGHV1-58*) (**Figure 1C and Figure S1C**). We also identified five immunoglobulin heavy diversity (*IGHD*) genes and one immunoglobulin heavy joining (*IGHJ*) gene. Within the light chain locus, we identified 11 functional immunoglobulin lambda variable (*IGLV*) genes and 77 *IGLV* pseudogenes, as well as one immunoglobulin lambda joining (*IGLJ*) gene (**Figure 1D**). The observation of multiple functional *IGLV* genes in the mallard duck was similar to that in the Muscovy duck^43^ and the Pekin duck^44^, but contrasted with the chicken, which possesses only a single functional *IGLV* gene^52^. Relatedly, our results revealed an expansion of *IGLV* pseudogenes in mallard ducks compared to the 33 identified in chickens^49^. Consistent with the evolutionary divergence between ducks and chickens, *IGHV1-1* and other *IGHV* pseudogenes in mallard ducks were phylogenetically distinct from those in chickens (**Figure 1E**). For example, the mallard duck *IGHV1-1* gene exhibited 77% sequence identity with the only functional *IGHV* gene (*IGHV1-1*) in chickens (**Figure S1D**). A similar observation could also be made for *IGLV* genes and pseudogenes (**Figure 1F and Figure S1E**). These findings establish a foundation for the molecular characterization of mallard duck antibody responses against influenza A virus.

**Figure 1.**
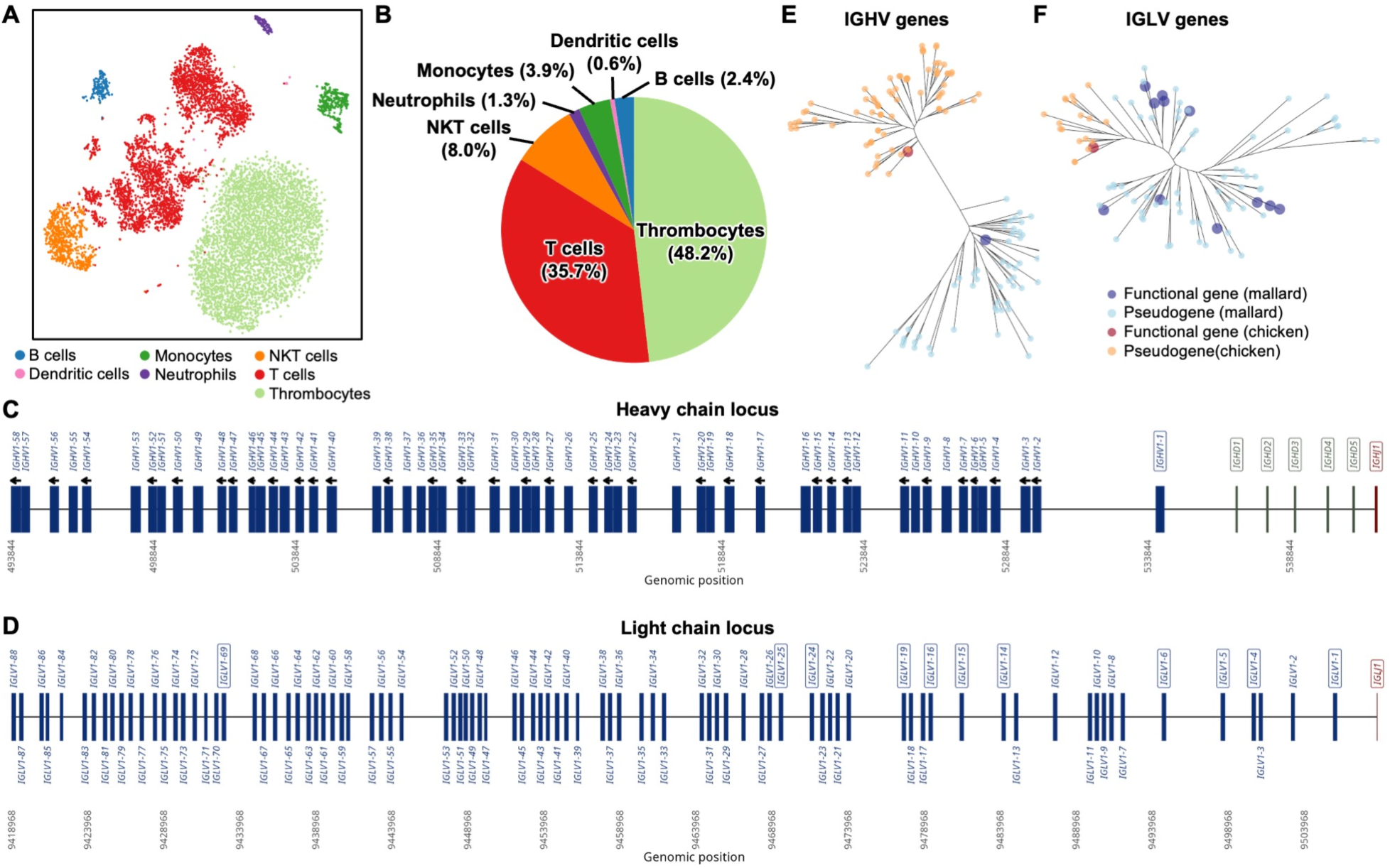
Annotation of mallard immunoglobulin germline genes. **(A)** Clustering of mallard PBMCs by T-distributed Stochastic Neighbor Embedding (t-SNE) analysis. Cell types were identified by the expression of selected gene markers (**see Methods**). **(B)** Distribution of cell types in mallard PBMCs. **(C-D)** The organization of the mallard **(C)** heavy chain and **(D)** light chain loci is shown, with *IGHV* (blue), *IGHD* (green), *IGHJ* (red), *IGLV* (blue), and *IGJV* (red) genes indicated. *IGHV* pseudogenes in reverse orientation relative to the functional *IGHV1-1* gene are indicated by black arrows. Names of functional genes are enclosed in boxes. **(E-F)** Phylogenetic comparison of **(E)** *IGHV* and **(F)** *IGLV* genes from chicken (orange) and mallard (blue). Enlarged tips denote functional genes.

### Isolation and functional characterization of duck HA antibodies

To study mallard duck antibody responses to influenza virus infection, six mallard ducks (group A) were sequentially infected with A/mallard/Alberta/328/2014 (H4N6) and A/mallard/Alberta/362/2017 (H3N8), and another six (group B) with A/mallard/Alberta/362/2017 (H3N8) and A/mallard/Alberta/566/85 (H6N2) (**Figure 2A**). These mallard ducks originated from the same laboratory colony as the one used for the above-mentioned immunoglobulin germline gene annotation. Both groups of ducks were able to generate antibody responses to the H3N8 HA, including some targeting the stem domain (**Figure 2B**). We also showed that these antibody responses have neutralization activity using an HA inhibition assay (**Figure 2C**) and a microneutralization assay (**Figure 2D**). Consistent with previous studies^30–33^, our results demonstrated that duck antibody responses to influenza HA could be elicited by experimental infection.

**Figure 2.**
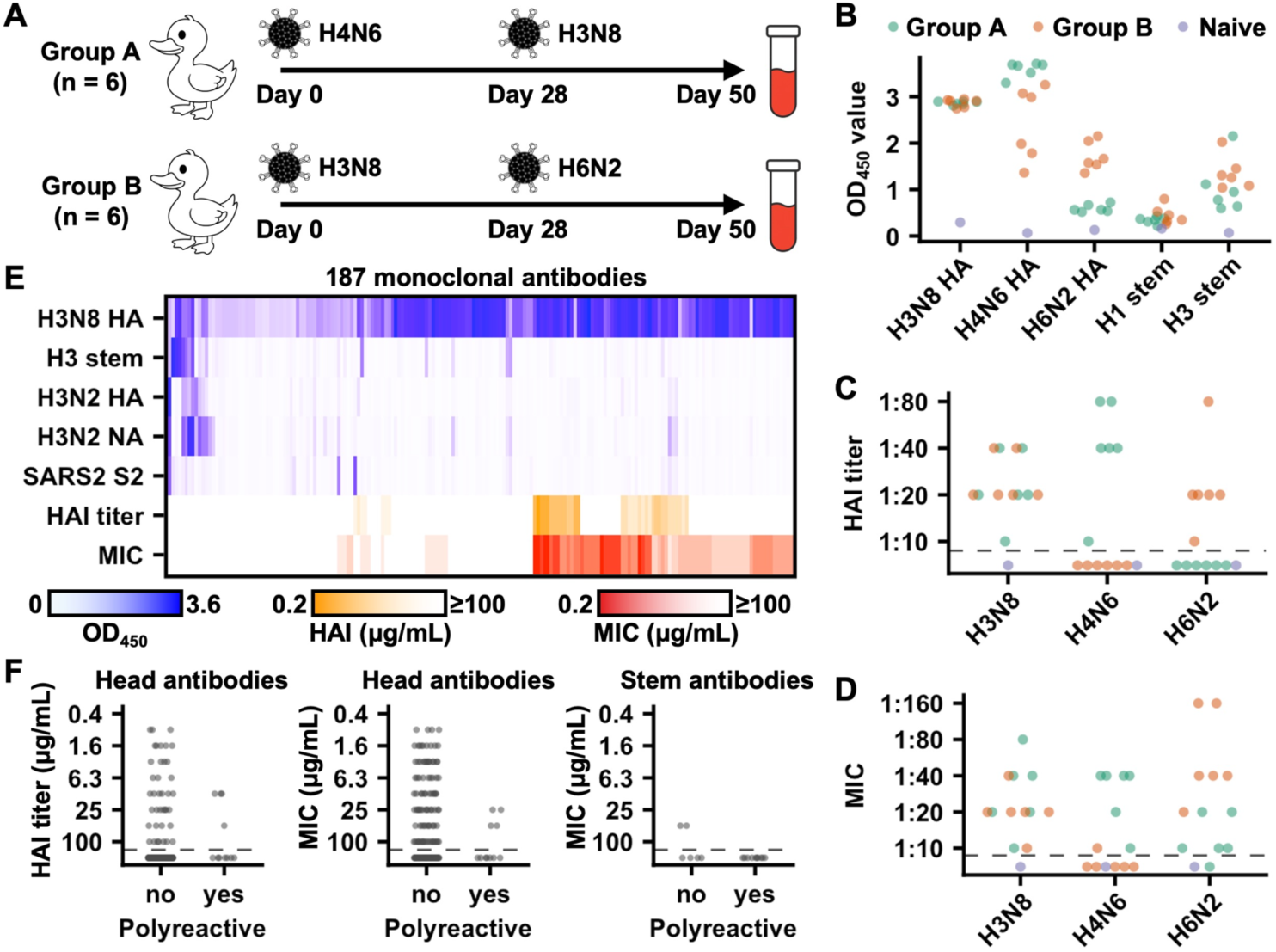
Isolation of monoclonal antibodies from experimentally infected mallard ducks. **(A)** Experimental design for mallard duck infection and blood collection. **(B)** Binding activity of plasma samples against the indicated antigens was measured by ELISA. **(C-D)** Neutralization activity of plasma samples against the indicated viruses was determined by **(C)** hemagglutination inhibition (HAI) assay and **(D)** microneutralization assay. **(B-D)** Green: group A mallard ducks. Orange: group B mallard ducks. Blue: naïve mallard ducks. MIC: minimum inhibitory concentration. Each datapoint represents the serum sample from one mallard duck. **(E)** Binding activity of 187 H3N8 HA antibodies to the indicated antigens and their neutralization activity as measured by HAI assay and microneutralization assay are shown as a heatmap. H3N8 HA: A/mallard/Alberta/362/2017 (H3N8) HA. H3 stem: a stabilized HA stem construct previously developed based on A/Finland/486/2004 (H3N2) HA^53^. H3N2 HA: A/Darwin/6/2021 (H3N2) HA. H3N2 NA: A/Moscow/10/1999 (H3N2) NA. SARS2 S2: SARS-CoV-2 S2 domain. Each column represents one antibody. Data represent the average of two highly consistent replicates. **(F)** Comparison of the neutralization activity against A/mallard/Alberta/362/2017 (H3N8) virus between polyreactive and non-polyreactive antibodies. Each datapoint represents one antibody. **(B-D and F)** Black dashed line represents the detection limit.

We further aimed to isolate monoclonal antibodies from the infected mallard ducks. A total of 2404 monoclonal antibodies were discovered by single-cell VDJ sequencing (scVDJ-seq) of PBMCs from group A and B mallard ducks sorted against H3N8 HA or an H3 stem construct previously developed based on A/Finland/486/2004 (H3N2) HA^53^ (**Figure S2A-B**). We also constructed a phage display antibody library from the PBMCs of the infected mallard ducks and screened it against H3N8 HA, resulting in additional 13 paired sequences of monoclonal antibodies. Among these 2417 mallard duck monoclonal antibodies (**Table S2**), 350 from diverse clonotypes were recombinantly expressed (**Figure S2C**), of which 187 showed reasonable binding activity (OD_450_ > 0.5) against H3N8 HA in ELISA, and a minor subset cross-reacted with A/Darwin/6/2021 (H3N2) HA (**Figure 2E and Table S3**). We further defined stem-binding antibodies as those with OD_450_ > 0.5 against H3 stem in ELISA. Among 172 validated H3N8 HA antibodies from scVDJ-seq of PBMCs sorted against H3N8 HA, only 14 (8.1%) were stem-binding (**Figure S2D**). Similarly, among 13 validated H3N8 HA antibodies from phage display screening, none bound to the stem domain. By contrast, the two validated H3N8 HA antibodies from scVDJ-seq of PBMCs sorted against H3 stem were both stem-binding. These observations showed that the HA head domain was immunodominant over the stem domain in ducks, consistent with findings in other animals, including humans^3,54,55^.

We also tested the polyreactivity of the 187 validated H3N8 HA antibodies. Here, we defined polyreactive antibodies as having an OD_450_ > 0.5 against A/Moscow/10/1999 (H3N2) neuraminidase (NA) or SARS-CoV-2 S2 domain in ELISA. Around 6% (11/171) of head-targeting antibodies and 63% (10/16) of stem-binding antibodies exhibited polyreactivity (**Figure 2E and Table S3**). Notably, the enrichment of polyreactivity in stem-binding antibodies has also been observed in humans^56^. Furthermore, all nine antibodies that cross-reacted with A/Darwin/6/2021 (H3N2) HA were polyreactive (**Figure 2E and Table S3**). Through HA inhibition assay and microneutralization assay, we observed that polyreactive antibodies tended to have weaker neutralization potency than non-polyreactivity antibodies (**Figure 2F**). For example, the median minimum inhibitory concentration (MIC) value of polyreactive head-targeting antibodies was >100 μg/mL, whereas that of non-polyreactive head-targeting antibodies was 50 μg/mL. Nevertheless, neutralization activity could still be detected for several polyreactive head-targeting antibodies. These results showed that polyreactivity was not uncommon in HA antibodies from mallard ducks, including some with neutralization activity.

Antibody polyreactivity is known to positively associate with CDR H3 length, hydrophobicity, and net charge^57–60^. However, the CDR H3 sequences of polyreactive and non-polyreactive head-targeting mallard duck antibodies had no significant difference in the hydrophobicity (p-value = 0.87, **Figure S2E**) and net charge (p-value = 0.94, **Figure S2F**). Additionally, the CDR H3 lengths of polyreactive head-targeting mallard duck antibodies were shorter than those of non-polyreactive ones, albeit with marginal statistical significance (p-value = 0.10, **Figure S2G**), which was opposite from previous findings on antibody polyreactivity^57–59^. Similarly, polyreactive and non-polyreactive stem-binding mallard duck antibodies showed no significant difference in their CDR H3 hydrophobicity (p-value = 0.21, **Figure S2E**), net charge (p-value = 0.90, **Figure S2F**), and length (p-value = 0.96, **Figure S2G**). Therefore, we concluded that other factors likely contributed to the polyreactivity of HA antibodies from mallard ducks.

### Polyreactive mallard duck HA antibodies bind to N-glycans

We hypothesized that polyreactive mallard duck HA antibodies bound to N-glycans, since the two antigens used for probing antibody polyreactivity, SARS-CoV-2 S2 domain and influenza virus NA (**Figure 2E and Table S3**), were both glycoproteins. Subsequently, we randomly selected seven polyreactive head-targeting antibodies and eight polyreactive stem-binding antibodies to test for binding to a panel of complex N-glycans by glycan array analysis (bottom panel in **Figure 3A, Figure S3A, and Table S4**) as well as to mannose-3 and mannose-9 glycans using biolayer interferometry (upper panel in **Figure 3A and Figure S3B**). Our results showed that 80% (12/15) of the selected polyreactive antibodies bound to at least one of the tested glycans. By contrast, none (0/9) of the selected non-polyreactive antibodies exhibited strong binding to the tested glycans **(Figure S3C-D)**.

**Figure 3.**
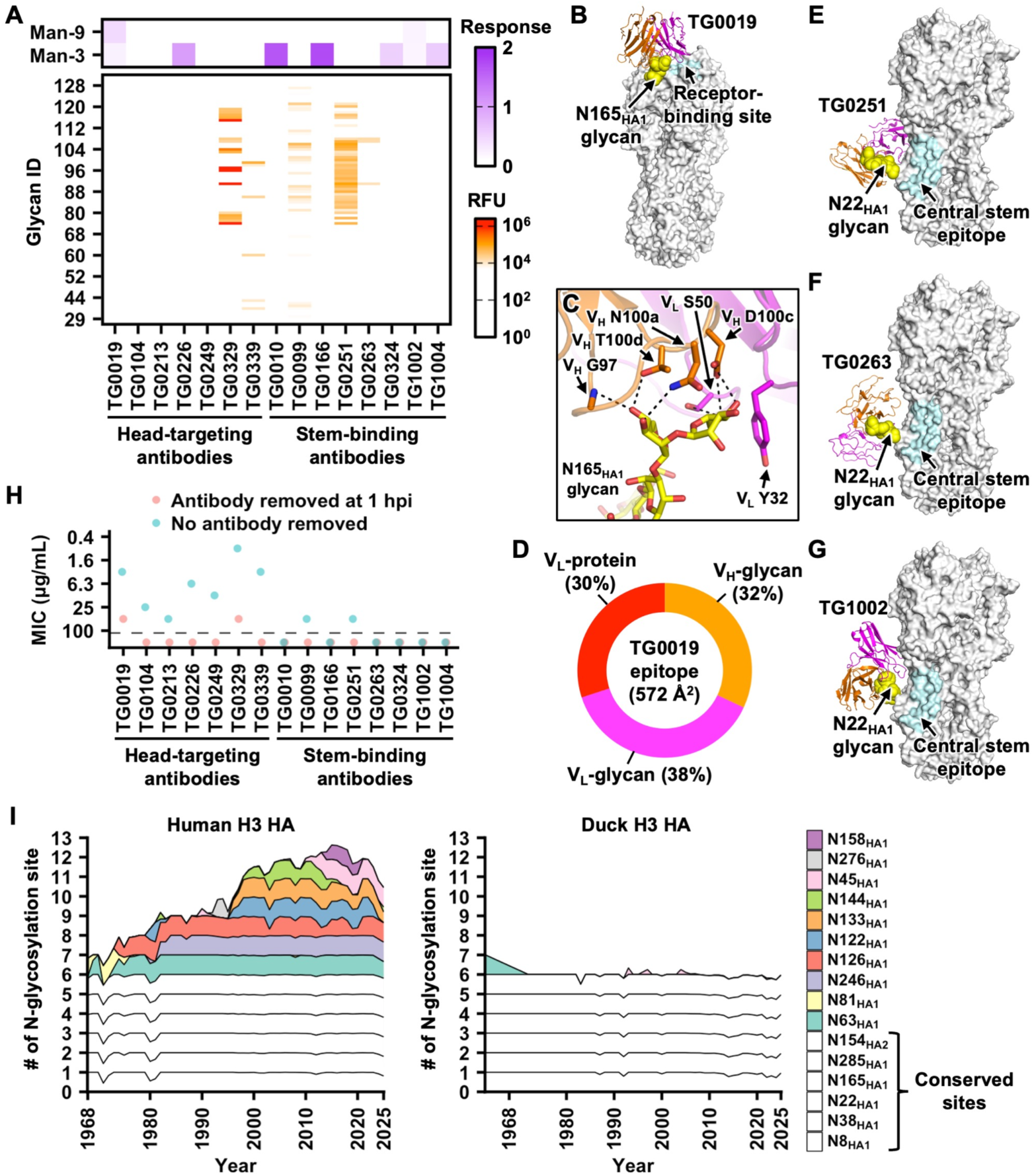
Characterization of polyreactive duck antibodies. **(A)** Glycan-binding activity of the indicated polyreactive antibodies was tested by glycan microarray (bottom panel, red) and biolayer interferometry (upper panel, purple). Man-9: mannose-9. Man-3: mannose-3. **(B)** Cryo-EM structure of TG0019 in complex with H3N8 HA. Orange: TG0019 heavy chain. Magenta: TG0019 light chain. White: HA. Yellow: N165_HA1_ glycan. Cyan: receptor-binding site. **(C)** Molecular interactions between TG0019 and N165_HA1_ glycan (yellow) are shown. Black dashed lines represent H-bonds. **(D)** The composition of TG0019 epitope is shown. V_L_-protein: interface between TG0019 light chain and the protein surface of HA. V_L_-glycan: interface between TG0019 light chain and N165_HA1_ glycan. V_H_-glycan: interface between TG0019 heavy chain and N165_HA1_ glycan. **(E-G)** Cryo-EM structures of **(E)** TG0251, **(F)** TG0263, and **(G)** TG1002 in complex with H3N8 HA. Orange: antibody heavy chain. Magenta: antibody light chain. White: HA. Yellow: N22_HA1_ glycan. Cyan: central stem epitope^3^. **(H)** Neutralization activity of the indicated polyreactive antibodies against A/mallard/Alberta/362/2017 (H3N8) virus is measured by microneutralization assay with (red) or without (blue) antibody removed at one-hour post-infection (hpi). MIC: minimum inhibitory concentration. Data from one of the two highly consistent replicates are shown. **(I)** The number of N-glycosylation sites on HA1 in human H3N2 strains since 1968 and in duck H3 strains since 1963.

To investigate how the glycan-binding ability of polyreactive antibodies contributes to HA interactions, we determined the cryo-EM structures of one polyreactive head-targeting antibody (TG0019) and four polyreactive stem-binding antibodies (TG0251, TG0263, TG1002, and TG1004) in complex with H3N8 HA to resolutions of 3.02 Å to 4.16 Å (**Figure S4 and Table S5**). TG0019, which bound more strongly to mannose-9 glycan and weakly to mannose-3 glycan (**Figure 3A and Figure S3B**), targeted the N165_HA1_ glycan in the head domain (**Figure 3B and Figure S3E**), which is highly conserved within H3 subtype^61^. TG0019 formed six H-bonds with two mannose moieties of the N165_HA1_ glycan, one of which belonged to the N-glycan core (**Figure 3C**). The N165_HA1_ glycan comprised 70% of the buried surface area of TG0019 epitope, with the remaining 30% attributed to light chain-protein interactions within the head domain (**Figure 3D**). Consistently, the binding activity of TG0019 to deglycosylated H3N8 HA was substantially reduced (**Figure S3F**). For the polyreactive stem-binding antibodies, TG0251, TG0263, and TG1002 all interacted with the N22_HA1_ glycan (**Figure 3E-G**), which is located adjacent to the central stem epitope, a common target for many human broadly neutralizing antibodies^3^. Lastly, while the density map for TG1004 was poorly resolved, it provided evidence that TG1004 targeted the N154_HA2_ glycan (**Figure S3G**). These analyses highlighted the critical role of glycan recognition in the binding of polyreactive mallard duck HA antibodies.

As described above, most polyreactive antibodies had minimal neutralization activity (**Figure 2F**). However, our microneutralization assay only measured the inhibition of virus entry, since antibodies were removed at one-hour post-infection, which was shorter than the approximately six-hour replication cycle of the influenza virus^62^. Recent studies have shown that some HA antibodies inhibit both virus entry and release^63,64^. To capture the inhibition of virus release by polyreactive antibodies, we performed an additional microneutralization assay that maintained antibodies in the medium for the duration of the experiment. In this microneutralization assay, virus growth inhibition was observed for all seven tested polyreactive head-targeting antibodies (**Figure 3H**). By contrast, six of the eight polyreactive stem-binding antibodies tested exhibited no viral growth inhibition, while the remaining two displayed weak activity with an MIC value of 50 μg/mL (**Figure 3H**). These results demonstrate the contribution of glycan-binding antibodies to the neutralizing activity of mallard duck antibody responses against HA.

To the best of our knowledge, none of the neutralizing HA antibodies isolated from influenza-infected humans have been shown to primarily target N-glycans. Consequently, mallard ducks appeared to have a greater propensity than humans to utilize HA N-glycans for neutralizing antibody responses. Such glycan-binding propensity suggested that adding N-glycans to H3 HA may not facilitate escape from duck antibody responses as effectively as for human responses^24,65^. This difference may also explain why human H3 HA evolved an extensive glycan shield^24,65^, whereas duck H3 HA remained relatively unshielded (**Figure 3I**).

### Recurrent engagement of the N22_HA1_ glycan by duck HA stem-binding antibodies

While most stem-binding antibodies are polyreactive (**Figure 2E and Table S3**), others, such as TG0331, are not (**Figure 4A**). Nonetheless, similar to those polyreactive stem-binding antibodies, TG0331 also broadly reacted against different HA subtypes (**Figure 4B**). Additionally, TG0331 exhibited weak neutralization activity (**Figure 4C**), comparable to some polyreactive stem-binding antibodies (**Figure 3H**). To understand how TG0331 targeted the stem domain, we determined a cryo-EM structure of TG0331 in complex with H3N8 HA to a resolution of 3.23 Å (**Figure 4D, Figure S5, and Table S5**).

**Figure 4.**
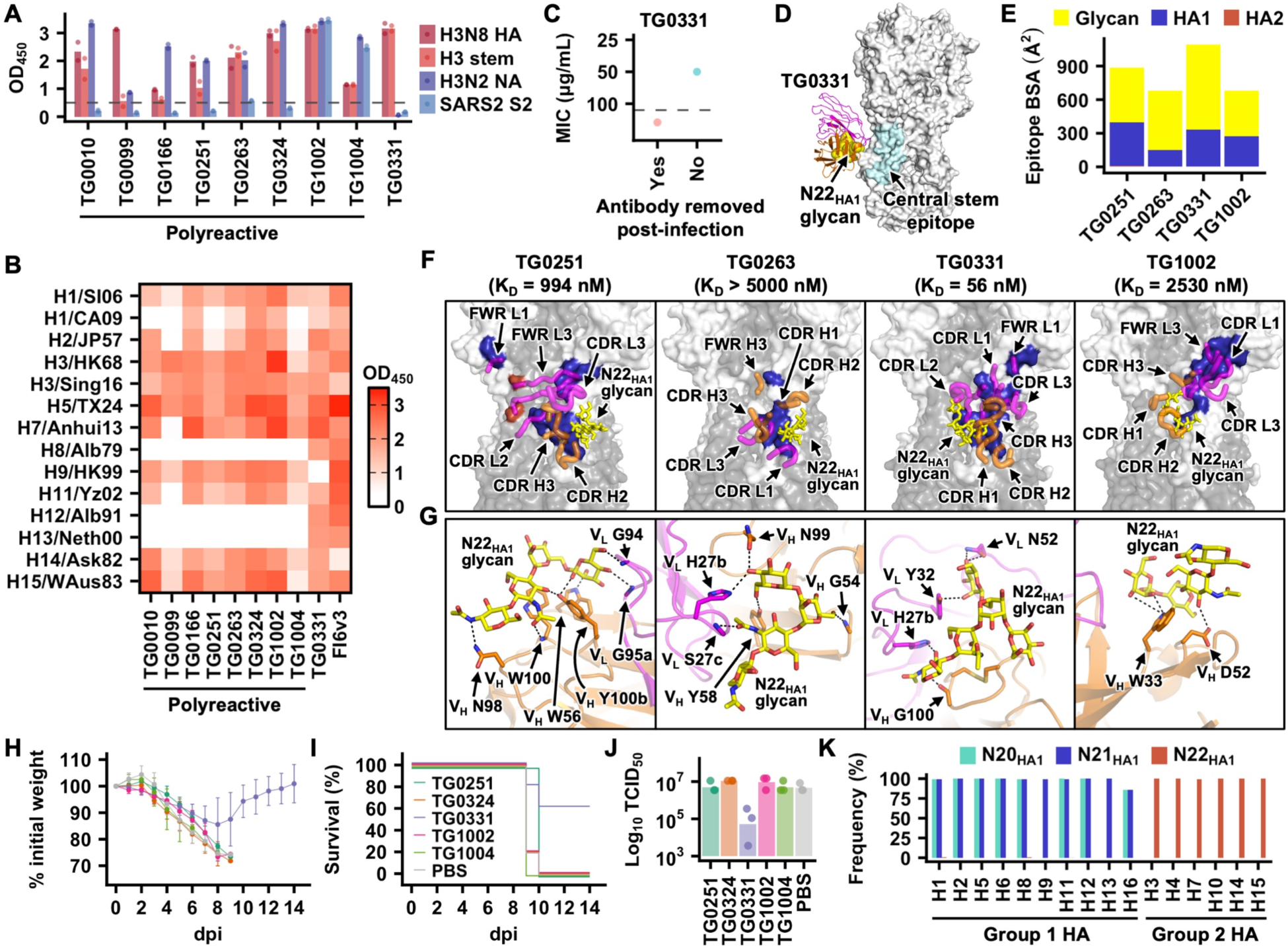
Characterization of stem-binding duck antibodies. (A-B) Binding activity of the indicated antibodies against the indicated antigens was measured by ELISA. **(A)** Each bar represents the average of two replicates. Each data point represents one replicate. H3N8 HA: A/mallard/Alberta/362/2017 (H3N8) HA. H3 stem: a stabilized HA stem construct previously developed based on A/Finland/486/2004 (H3N2) HA^53^. H3N2 NA: A/Moscow/10/1999 (H3N2) NA. SARS2 S2: SARS-CoV-2 S2 domain. **(B)** The average of two replicates, which are highly consistent, is shown. H1/SI06: A/Solomon Island/3/2006 (H1N1). H1/CA09: A/California/07/2009 (H1N1). H2/JP57: A/Japan/305/1957 (H2N2). H3/HK68: A/Hong Kong/1968 (H3N2). H3/Sing16: A/Singapore/INFIMH-16-0019/2016 (H3N2). H5/TX24: A/cattle/Texas/56283/2024 (H5N1). H7/Anhui13: A/Anhui/1/2013 (H7N9). H8/Alb79: A/pintail duck/Alberta/114/1979 (H8N4). H9/HK99: A/Hong Kong/1073/1999 (H9N2). H11/Yz02: A/duck/Yangzhou/906/2002 (H11N2). H12/Alb91: A/green-winged teal/ALB/199/1991 (H12N5). H13/Neth00: A/black-headed gull/Netherlands/1/2000 (H13N8). H14/Ask82: A/mallard/Astrakhan/263/1982 (H14N5). H15/WAus83: A/Australian shelduck/Western Australia/1756/1983 (H15N2). **(C)** Neutralization activity of TG0331 against A/mallard/Alberta/362/2017 (H3N8) virus was measured by microneutralization assay with (red) or without (blue) antibody removed at one-hour post-infection (hpi). MIC: minimum inhibitory concentration. Data from one of the two highly consistent replicates are shown. **(D)** The cryo-EM structure of TG0331 in complex with H3N8 HA. Orange: antibody heavy chain. Magenta: antibody light chain. White: HA. Yellow: N22_HA1_ glycan. Cyan: central stem epitope^3^. **(E)** The composition of TG0331 epitope is shown. HA1: protein surface of HA1. HA2: protein surface of HA2. Glycan: N22_HA1_ glycan. **(F)** Epitopes of the indicated antibodies are in blue. Their K_D_ values are indicated in the bracket. Yellow: N22_HA1_ glycan. **(G)** Interactions between the indicated antibodies and N22_HA1_ glycan (yellow) are shown. Black dashed lines represent H-bonds. **(H-J)** Female BALB/c mice at 6 weeks old were injected intraperitoneally with 5 mg kg^-1^ of the indicated antibody at 4 h prior to challenge with 5× LD_50_ of mouse-adapted H3N2 A/Philippines/2/1982 (X-79, 6:2 A/PR/8/34 reassortant) virus. **(H)** The mean percentage of body weight change post-infection is shown (n = 5). Error bars indicate standard deviation. The humane endpoint was defined as a weight loss of 25% from initial weight on day 0. Different colors represent different antibodies as indicated in panel I. **(I)** Kaplan-Meier survival curves are shown (n = 5). **(J)** Lung viral titers on day 3 post-infection are shown (n = 3). **(K)** Occurrence frequency of N20_HA1_ glycan, N21_HA1_ glycan, and N22_HA1_ glycan in different HA subtypes.

Although TG0331 was not classified as a polyreactive antibody (**Figure 2E**, **Figure 4A, and Table S3**) and had no detectable binding activity to the tested glycans above (**Figure S3C-D**), it also engaged the N22_HA1_ glycan as observed for TG0251, TG0263, and TG1002 (**Figure 3E-G and Figure 4D**). For all four antibodies, the light chain dominated the binding interface relative to the heavy chain, accounting for 52% to 59% of their paratopes (**Figure S6A**). Moreover, 55% to 78% of their epitopes were attributed to N22_HA1_ glycan (**Figure 4E**). Consistently, their binding activity to deglycosylated H3N8 HA was substantially reduced (**Figure S6B**). The N22_HA1_ glycan was positioned to the right side of TG0251 and TG0263, but to the left side of TG0331 and TG1002 (**Figure 4F and Figure S6C-F**), showcasing the diverse approach angles of mallard duck antibodies to the N22_HA1_ glycan. All four antibodies interacted with the N-glycan core, Manα1–3(Manα1–6)Manβ1-4GlcNAcβ1–4GlcNAcβ1, albeit via distinct H-bond networks (**Figure 4G**). Furthermore, the heavy chains of TG0251, TG0331, and TG1002 interacted with the lower part of the epitope, while that of TG0263 interacted with the upper part (**Figure 3E-G**, **Figure 4D, and Figure 4F**). The binding affinity of TG0251 and TG0331 in Fab format was in the nM range (58 nM and 994 nM, respectively), whereas that of TG0263 and TG1002 was in the μM range (**Figure S6G**). The stronger binding affinity of TG0251 and TG0331 correlated with their larger epitope surface area (**Figure 4E**) and their detectable neutralization activity (**Figure 3H and Figure 4C**). Similarly, only TG0331, but none of the other tested stem-binding antibodies with weaker affinity (TG0251, TG0324, TG1002, and TG1004), conferred protection against lethal H3N2 challenge in a mouse model, as demonstrated by weight loss (**Figure 4H**), survival (**Figure 4I**), and viral lung titer at day 3 post-infection (**Figure 4J**). Together, our analyses revealed the N22_HA1_ glycan as a common target of stem-binding mallard duck antibodies albeit with different binding modes.

Previous studies on H7 subtype demonstrate the importance of the N22_HA1_ glycan (N12_HA1_ glycan in H7 numbering) for HA stability, and, consequently, viral replication^66,67^. Consistently, the N22_HA1_ glycosylation site is highly conserved among group 2 HAs, including H3, H4, H7, H10, H14, and H15 (**Figure 4K**). By contrast, group 1 HAs lack the N22_HA1_ glycosylation site, but possess an N-glycosylation site at N21_HA1_. Nevertheless, the spatial location of N21_HA1_ in group 1 HA was similar to that of N22_HA1_ in group 2 HAs (**Figure S6H**), which could explain the cross-reactivity of TG0251, TG0263, TG0331, and TG1002 to group 1 HAs (**Figure 4B**).

### Duck antibody responses engage diverse HA head epitopes

As shown above, non-polyreactive head-targeting antibodies typically had more potent neutralization activity than polyreactive or stem-binding antibodies (**Figure 1F**). To identify the HA head epitopes that are targeted by mallard duck antibodies, we determined cryo-EM structures of 12 randomly selected non-polyreactive head-targeting antibodies to resolutions of 2.34 Å to 3.33 Å (**Figure 5, Figure S5, Figure S7, Figure S8, Figure S9A and Table S5**). These antibodies included eight from scVDJ-seq of PBMCs sorted against H3N8 HA and three from phage display screening. Most of these non-polyreactive head-targeting antibodies primarily used the heavy chain for binding, although three exhibited light-chain-dominant binding (**Figure S9B and Figure S10**). Consistent with the observation that their binding activity is maintained even after HA deglycosylation (**Figure S9C**), none of these 12 antibodies engaged N-glycans. Nonetheless, their epitopes were highly diverse, with each epitope spanning multiple major antigenic sites (**Figure 5A and Figure S9D**). For example, the TG0009 epitope overlapped with sites C, D, and E, whereas that of TG0253 overlapped with sites A, B, and D (**Figure S9D**). Collectively, they encompassed nearly the entire HA head domain with a relatively uniform distribution (**Figure 5A-B**). Except for site C, which is located on the lower part of the head domain and was targeted by only three of the 12 antibodies, sites A, B, D, and E were targeted by nine to 11 antibodies (**Figure 5C**). As a result, the mallard duck antibody responses displayed a more balanced immunodominance hierarchy than human antibody responses against H3 HA, which mainly focus on sites A and B^68–71^.

**Figure 5.**
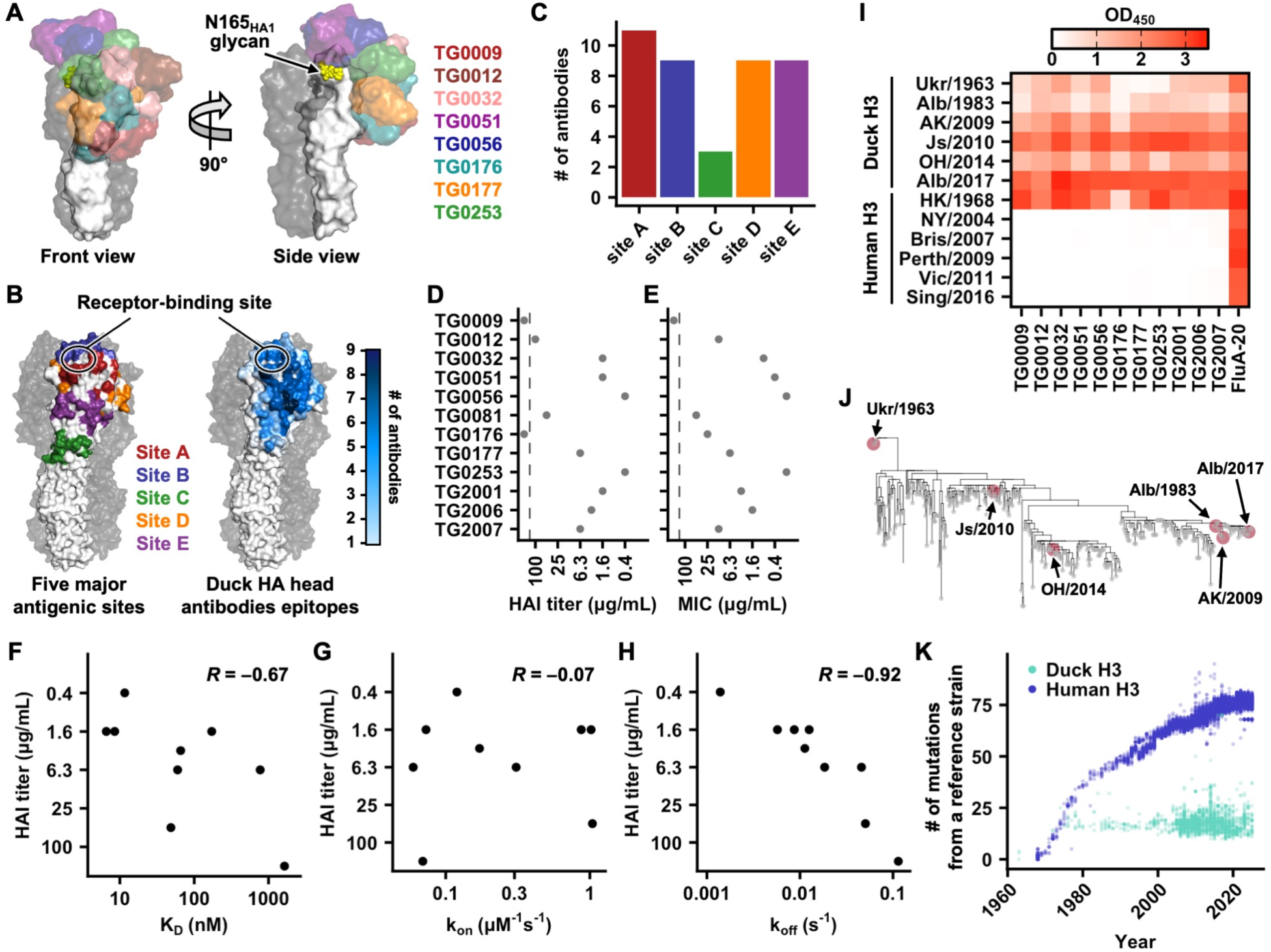
Characterization of head-targeting duck antibodies. **(A)** Structural overlays of representative head-targeting duck antibodies are shown. **(B)** Locations of the five major antigenic sites on H3 HA^19,21,135^ are shown on the left. HA head surfaces that are targeted by duck antibodies are shown on the right, with residues colored according to the frequency of being targeted by duck antibodies (out of 12). **(C)** Among 12 head-targeting antibodies with cryo-EM structures determined, the number of antibodies targeting each of the five major antigenic sites^19,21,135^ is shown. **(D-E)** Neutralization activity of those head-targeting duck antibodies with cryo-EM structures determined against A/mallard/Alberta/362/2017 (H3N8) virus was measured using **(D)** HAI assay and **(E)** microneutralization assay. MIC: minimum inhibitory concentration. Data from one of the two highly consistent replicates are shown. **(F-H)** The relationship between HAI titers and binding kinetics, including **(F)** K_D_, **(G)** on-rate (k_on_), and **(H)** off-rate (k_off_) was analyzed for nine head-targeting duck antibodies. Binding kinetics were measured using Fabs. **(I)** Binding activity of the indicated antibodies against the indicated H3 HAs was measured by ELISA. Data represent the average of two highly consistent replicates. **(J)** Phylogenetic tree of duck H3 HAs. Red: HAs that were included in our binding analysis in panel I. **(K)** The number of amino acid mutations in the HA ectodomain is shown for human H3 strains (blue) relative to the reference strain A/Hong Kong/1/1968 (H3N2) (blue) and for duck H3 strains (cyan) relative to A/duck/Ukraine/1/1963 (H3N8). Each data point represents one strain.

The 12 non-polyreactive head-targeting antibodies had a broad range of neutralization activity as measured by hemagglutination inhibition (HAI) and microneutralization assays (**Figure 5D-E and Table S3**). For example, HAI activity of TG0056 and TG0253 was observed at concentrations as low as 0.4 μg/mL, whereas no HAI activity was detected for TG0009 and TG0176, even at 100 μg/mL (**Figure 5D**). We further expressed and purified nine of these head-targeting antibodies in Fab format for binding kinetics measurements using biolayer interferometry (**Figure S9E-F**). Their neutralization activity correlated strongly with the binding off-rate (Pearson correlation coefficient = −0.92) and moderately with the K_D_ (Pearson correlation coefficient = −0.67), but showed no correlation with the on-rate (**Figure 5F-H and Figure S9G-I**). This observation indicated that binding off-rate was the primary determinant of neutralization activity for head-targeting antibodies across different epitopes.

Compared to human H3 HA, duck H3 HA has a relatively stable antigenicity^25–28^. Consistently, most head-targeting antibodies exhibited broad cross-reactivity with all tested HAs from duck H3 strains (**Figure 5I**), which spanned a period from 1963 to 2017 and were phylogenetically distinct (**Figure 5J**). By contrast, these head-targeting antibodies only bound to the human H3 HA from the ancestral human H3N2 strain A/Hong Kong1/1968, but not those from more recent strains (**Figure 5I**). These results corroborated the observation that duck H3 HA accumulated amino acid mutations at a lower rate than human H3 HA (**Figure 5K**)^29^. Nevertheless, the antigenicity of duck H3 HA still evolves over time, albeit very slowly. For example, TG0176 and TG0177 bound strongly to A/mallard/Alberta/362/2017 (H3N8) HA but very weakly to A/duck/Ukraine/1/1963 (H3N8) HA (**Figure 5I**), likely due to the amino acid mutation at HA1 residue 63. While the D63_HA1_ side chain of A/mallard/Alberta/362/2017 (H3N8) HA formed H-bonds with TG0176 and TG0177 (**Figure S10**), such H-bonds would be absent in A/duck/Ukraine/1/1963 (H3N8) HA, which contains A63_HA1_ instead (**Figure S9J**). Since the balanced targeting of multiple epitopes by mallard duck antibody responses would create a high genetic barrier that requires multiple concurrent mutations for escape, as shown in human antibody responses to measles virus^72,73^, it may explain, at least in part, the antigenic stability of duck H3 HA.

### A convergent CDR H3- and light chain-independent binding mode of four antibodies

Among the 12 non-polyreactive head-targeting antibodies for which we determined the structures, one from scVDJ-seq (TG0012) and three from phage display screening (TG2001, TG2006, and TG2007), targeted an almost identical epitope in the HA head domain using highly similar heavy chain-only binding modes (**Figure 6A**). The CDR H3 sequences of these four antibodies were diverse and had no contact with the HA (**Figure 6B and Figure S11D**), although antibody binding independent of CDR H3 is rare^74^. Instead, TG0012, TG2001, TG2006, and TG2007 mainly relied on CDR H2 and framework region (FWR) H3 for binding to HA via extensive H-bond networks (**Figure 6C and Figure S11A-C**).

**Figure 6.**
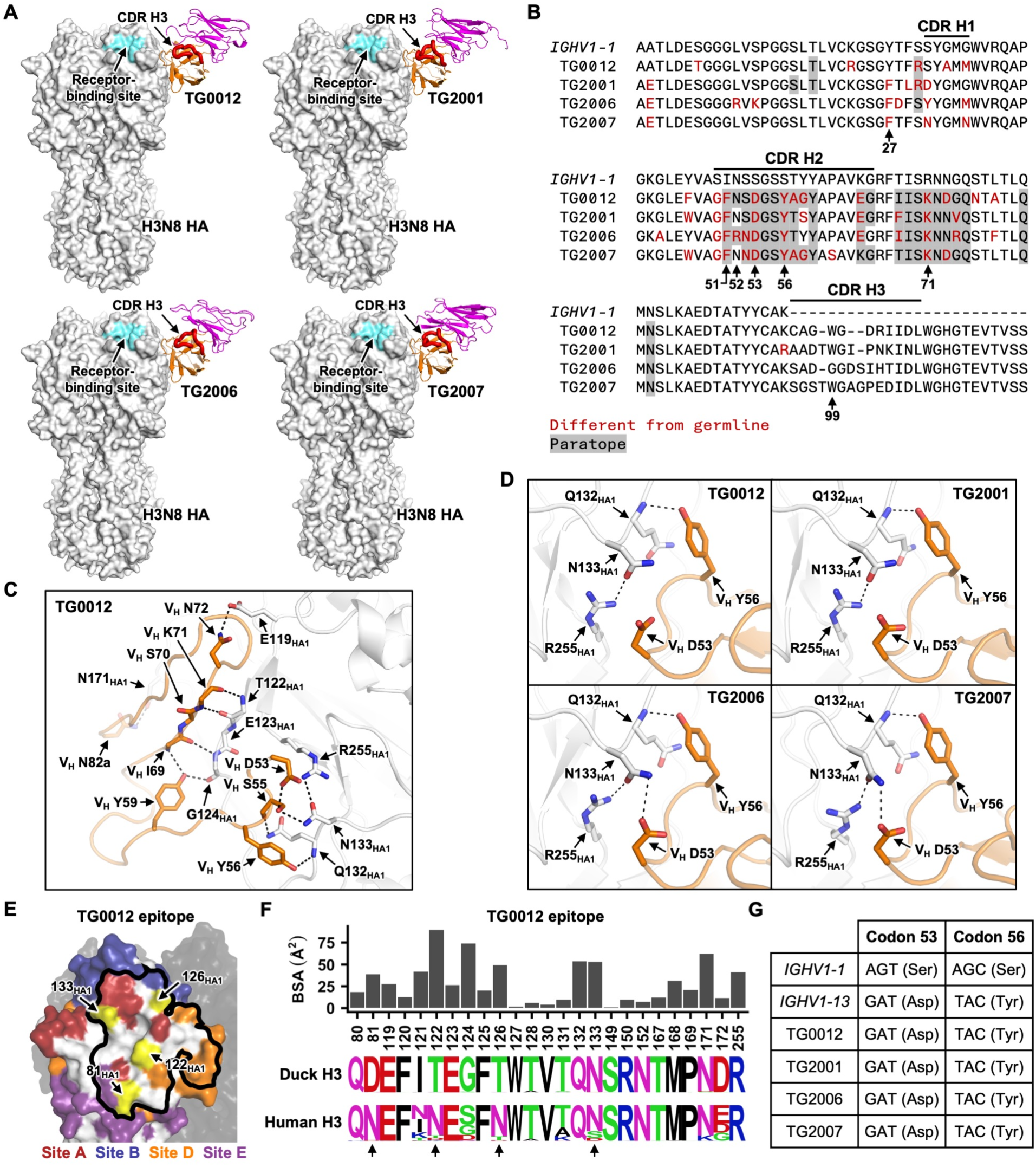
A convergent antibody binding mode without the involvement of CDR H3 and light chain. **(A)** Cryo-EM structures of TG0012, TG2001, TG2006, and TG2007 in complex with H3N8 HA. Orange: antibody heavy chain. Magenta: antibody light chain. White: HA. Cyan: receptor-binding site. **(B)** Alignment of TG0012, TG2001, TG2006, and TG2007 heavy chain sequences with mallard *IGHV1-1* germline gene. Amino acid differences from *IGHV1-1* germline gene are shown red, whereas paratope residues are shaded in gray. Residues of interest are indicated. **(C)** Key residues at the interface between TG0012 and HA are shown. White: HA. Orange: TG0012 heavy chain. Black dashed lines represent H-bonds or electrostatic interactions. **(D)** The interactions of HA with V_H_ D53 and V_H_ Y56 in TG0012, TG2001, TG2006, and TG2007 are shown. **(E)** TG0012 epitope is outlined on one HA protomer with the major antigenic sites indicated. The other two HA protomers are shown in black. N-glycosylation sites with high occurrence frequency in human H3 strains are shown in yellow. **(F)** The bar chart indicates the buried surface area (BSA) of each residue in the TG0012 epitope upon binding. The sequence logos represent the sequence diversity of each epitope residue in duck H3 strains from 1963 to 2025, as well as human H3 strains from 1968 to 2025. N-glycosylation sites present in human H3 strains but not duck H3 strains are indicated by arrows. **(G)** Codon usage at V_H_ residues 53 and 56 of *IGHV1-1*, *IGHV1-13*, TG0012, TG2001, TG2006, and TG2007. The encoded amino acids are shown in brackets.

TG0012, TG2001, TG2006, and TG2007 shared several amino acid mutations that were different from the functional germline gene *IGHV1-1* sequence, including V_H_ F51, V_H_ D53, V_H_ Y56, and V_H_ K71 (**Figure 6B**). In all four antibodies, V_H_ D53 formed favorable electrostatic interactions with R255_HA1_, whereas V_H_ Y56 H-bonded with the backbone amide nitrogen of Q132_HA1_ (**Figure 6D**). In TG2006 and TG2007, V_H_ D53 also H-bonded with N133_HA1_. These interactions would not exist if V_H_ residues 53 and 56 were encoded by the functional germline gene *IGHV1-1*, which had Ser at these two residues (**Figure 6B**). Our sequence analysis showed that both V_H_ S53D and V_H_ S56Y mutations in TG0012, TG2001, TG2006, and TG2007 required two nucleotide changes from the *IGHV1-1* (**Figure 6G**). *IGHV1-13* was the only pseudogene that encoded both V_H_ D53 and V_H_ Y56, with a codon usage identical to those of TG0012, TG2001, TG2006, and TG2007 (**Figure 6G**). Thus, TG0012, TG2001, TG2006, and TG2007 likely acquired V_H_ D53 and V_H_ Y56 through gene conversion from *IGHV1-13* rather than somatic hypermutation. Similarly, V_H_ K71, which formed intramolecular interactions in the protein core (**Figure S11E**), was present in multiple pseudogenes (**Figure S1C**). By contrast, none of the pseudogenes had V_H_ F51, which therefore must have arisen due to somatic hypermutation. These observations showed that gene conversion and somatic hypermutation, rather than VDJ recombination, were critical for the binding of TG0012, TG2001, TG2006, and TG2007 to HA.

The epitope of TG0012, TG2001, TG2006, and TG2007, which partly overlapped with antigenic sites A, B, D, and E (**Figure 6E and Figure S11F**), was highly conserved among H3 strains circulating in ducks but not among those in humans (**Figure 6F and Figure S11G-I**). This finding was consistent with the broad reactivity of these four antibodies against diverse duck H3 HAs (**Figure 5I-J**). Within this epitope, human H3 strains contained four N-glycosylation sites that were absent in duck H3 strains, namely N81_HA1_, N122_HA1_, N126 _HA1_, and N133_HA1_ (**Figure 3I**)^23^. The lack of binding of TG0012, TG2001, TG2006, and TG2007 to most human H3 HAs tested was likely due to the presence of one or more of these N-glycosylation sites, which may sterically hinder antibody access (**Figure 5I**). However, these four antibodies could still bind to ancestral human H3N2 strain A/Hong Kong1/1968, despite the presence of the N81_HA1_ glycan.

### A head-targeting antibody utilizing an N-glycan as a decoy receptor

TG0081 was another non-polyreactive head-targeting antibody with uncommon structural features, owing to an N-glycosylation site at residue V_H_ N100a in the CDR H3 (**Figure 7A**). This N-glycosylation site was encoded by N-nucleotide addition (**Figure S12A**). While the protein portion of TG0081 targeted an epitope that spanned major antigenic sites A, B, D, and E (**Figure 7B-C**), its V_H_ N100a glycan engaged the receptor-binding site of an adjacent HA protomer via a terminal sialic acid moiety (**Figure 7B and Figure S12B**). In fact, the V_H_ N100a glycan was a major contributor to the binding interface, accounting for 52% of the paratope (**Figure 7D**). The binding of the protein portion of TG0081 to HA involved multiple H-bonds and a π-π interaction between V_L_ Y31 and Q132_HA1_ (**Figure 7E**). At the same time, the V_H_ N100a glycan extended from CDR H3 to form two H-bonds with the N165_HA1_ glycan of the same HA protomer before reaching the receptor-binding site of the adjacent protomer (**Figure 7F**). The position of the V_H_ N100a glycan was stabilized by the intramolecular interactions formed by the V_H_ N100a side chain, which H-bonds with the V_H_ S99 side chain and the V_H_ W100b backbone amide nitrogen, alongside a π-π interaction with the V_H_ Y56 side chain. V_H_ Y56 was likely acquired through gene conversion from *IGHV1-23*, as the TG0081 amino acid sequence from V_H_ residues 47 to 61 was identical to *IGHV1-23* but differed from *IGHV1-1* by eight amino acids (**Figure S12C**).

**Figure 7.**
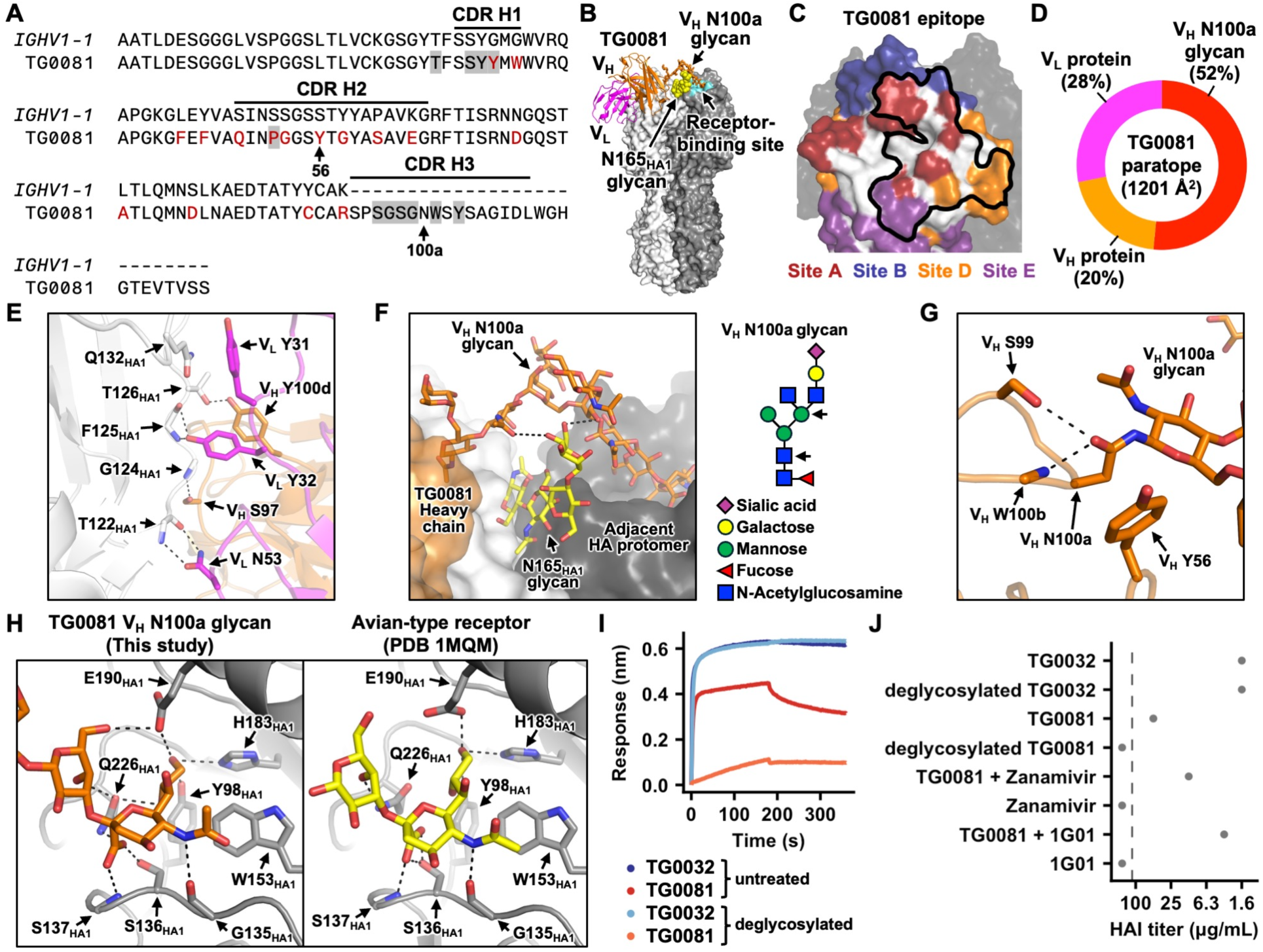
Characterization of TG0081 and the importance of an N-glycan on its CDR H3. **(A)** Alignment of TG0081 heavy chain sequence with mallard *IGHV1-1* germline gene. Amino acid differences from *IGHV1-1* germline gene are shown red, whereas paratope residues are shaded in gray. **(B)** Cryo-EM structure of TG0081 in complex with H3N8 HA. Orange: antibody heavy chain. Magenta: antibody light chain. White: HA. Cyan: receptor-binding site. Cyan: receptor-binding site. **(C)** Epitope targeted by the protein portion of TG0081 is outlined on one HA protomer with the major antigenic sites indicated. The other two HA protomers are shown in black. **(D)** The composition of TG0081 paratope is shown. V_H_ protein: protein surface of heavy chain. V_L_ protein: protein surface of light chain. V_H_ N100a glycan: the N-glycan at V_H_ residue N100a. **(E)** Key residues at the interface between TG0081 and HA are shown. White: HA. Orange: TG0081 heavy chain. Magenta: TG0081 light chain. **(F)** Interactions between the TG0081 V_H_ N100a glycan (orange) and the N165_HA1_ glycan (yellow) are shown. The HA protomer interacting with the protein surface of TG0081 is in white, whereas that interacting with the TG0081 V_H_ N100a glycan is in black. Glycan diagram for the resolved structure of TG0081 V_H_ N100a glycan is shown on the right according to the Symbol Nomenclature for Glycans recommended by the NLM ^136,137^. Arrows indicate the moieties that interact with the N165_HA1_ glycan. **(G)** Intramolecular interaction involving TG0081 V_H_ N100a side chain is shown. **(H)** Structural comparison between the binding of TG0081 V_H_ N100a glycan and an avian-type receptor analog (PDB 1MQM)^138^ to the H3 HA receptor-binding site. **(E-H)** Black dashed lines represent H-bonds. **(I)** The binding of H3N8 HA to TG0032 IgG and TG0081 IgG, either with or without deglycosylation, was measured using biolayer interferometry. **(J)** The neutralization activity of TG0032 IgG and TG0081 IgG, either with or without deglycosylation, against A/mallard/Alberta/362/2017 (H3N8) virus was measured using HAI assay. The effects of zanamivir and 1G01 on TG0081 neutralization activity were also tested.

The terminal sialic acid moiety of TG0081 V_H_ N100a glycan was α2-3 linked, matching the linkage found in avian-type receptors for influenza A virus^3^. Consistently, the interaction of HA receptor-binding site with the terminal sialic acid moiety of TG0081 V_H_ N100a glycan was similar to its interactions with the avian-type receptor (**Figure 7F**). Upon deglycosylation, the binding activity of TG0081 was markedly reduced (**Figure 7I**), and its neutralizing activity became undetectable (**Figure 7J**), substantiating the importance of the V_H_ N100a glycan for binding. As a control, deglycosylation of TG0032, another head-targeting antibody (**Figure 5A**), had no effect on its binding or neutralizing activity. Together, our data demonstrated that the V_H_ N100a glycan allowed TG0081 to partially function as a decoy receptor. To the best of our knowledge, this antibody binding mechanism has not been observed in any previously described influenza HA antibodies, despite the presence of N-glycans in the Fab region of around 14% of human antibodies^75^.

Since influenza NA on the viral surface cleaves sialic acid^76^, we postulated that it could also remove the sialic acid from the V_H_ N100a glycan and that the neutralization potency of TG0081 could be enhanced by inhibiting the NA activity. Subsequently, we tested whether the neutralization potency of TG0081 would be affected by zanamivir, a small-molecule NA inhibitor, or 1G01, an antibody targeting the NA active site^77^. While zanamivir or 1G01 alone had no HAI activity, their presence could increase the HAI activity of TG0081 (**Figure 7J**). For example, the HAI titer of TG0081 improved more than 10-fold from 50 μg/mL to 3.1 μg/mL, when 100 μg/mL of 1G01 was added. This observation not only illustrated that influenza NA activity can negatively impact the potency of TG0081, but also suggested a synergistic effect between TG0081 and NA immunity.

## DISCUSSION

As the natural reservoir, waterfowl species such as mallard ducks are central to the ecology of influenza A virus^6^. However, the molecular characteristics of immune pressures encountered by influenza A virus in the natural reservoir have remained largely unclear, especially since the antigenicity of at least some subtypes can be relatively stable despite widespread immunity in waterfowl^25–28,35–38,51^. By using the H3 subtype as a model, we performed an in-depth molecular investigation of mallard duck antibody responses against influenza virus infection, spanning from immunoglobulin germline discovery to the isolation and characterization of monoclonal antibodies. Our work not only reveals the uniqueness of mallard duck antibody responses, but also offers important mechanistic insights into the antigenic stability of influenza A virus in the natural reservoir.

A notable finding in this study is the identification of multiple neutralizing antibodies that predominantly target N-glycans. Although thousands of neutralizing antibodies have been isolated from human influenza virus infections or vaccinations^78^, to the best of our knowledge, none have shown strong dependence on N-glycans for binding. The propensity of mallard ducks to produce glycan-binding antibodies helps explain the lack of N-glycosylation site accumulation in influenza HA within the natural reservoir. This propensity is likely encoded by the duck immunoglobulin light chain genes, since all five glycan-binding antibodies in our structural analysis (TG0019, TG0251, TG0263, TG0331, and TG1002) rely predominantly on the light chain, rather than heavy chain, for binding. It is also possible that this propensity is associated with the expansion of functional *IGLV* genes and *IGLV* pseudogenes in ducks relative to chickens. Nonetheless, we have minimal, if any, molecular understanding of chicken antibody responses to the influenza HA. Comparative studies of various avian antibody responses are necessary to clarify the immunological relevance of the expansion of *IGLV* genes in ducks.

The minimal glycan shielding in duck H3 HAs leaves most of the head domain exposed to antibody targeting, potentially contributing to the relatively balanced immunodominance hierarchy of duck antibody responses across different antigenic sites. By contrast, human antibody responses to human H3 HA mainly focus on sites A and B^68–71^, a tendency that is reinforced by the progressive accumulation of N-glycosylation sites in other regions of the head domain^24,65^. For instance, in a previous study of 33 H3 HA antibodies from human vaccinees, only 7 (21%) recognized epitopes outside of sites A and B^71^. The human antibody responses are so focused that even a single H3 HA mutation can often drastically reduce, if not abolish, the neutralization activity of human sera^79,80^. We postulate that a similar reduction in the neutralization activity of duck sera against H3 HA would require multiple concurrent mutations, given the balanced immunodominance hierarchy. Consequently, the selective advantage of a single mutation conferred by duck immune pressure is likely minimal and outweighed by the associated replication fitness cost. This notion is parallel to the observation that the antigenic evolution of measles virus is constrained by having multiple co-dominant epitopes^72,73^. As a result, the combination of a balanced immunodominance hierarchy and a propensity for generating glycan-binding antibodies in ducks would synergistically minimize the antigenic drift of influenza A virus in the natural reservoir.

Another highlight of this study is the CDR H3-independent binding mechanism exhibited by antibodies TG0012, TG2001, TG2006, and TG2007, which share the same epitope that is conserved in duck H3 HA. Without a strict requirement for CDR H3 sequences or specific VDJ recombination, this binding mode would allow ducks to rapidly mount neutralizing responses against the H3 subtype. This class of antibodies could be a result of the coevolution between dabbling ducks and influenza A virus, as the co-occurrence of the two key binding residues, V_H_ D53 and V_H_ Y56, is unique to mallard *IGHV1-13* but absent from both the functional gene *IGHV1-1* and all pseudogenes in chickens^49^. It also exemplifies the importance of gene conversion in generating mallard duck HA antibodies. Notably, this CDR H3-independent binding mechanism is reminiscent of the interaction between a human IgM antibody and a superantigen from *Staphylococcus aureus*^81^, which is mediated exclusively by heavy-chain framework residues. Nevertheless, whether duck antibodies targeting other HA subtypes also utilize CDR H3-independent binding mechanism is an open question.

Although this study establishes the molecular basis of the interactions between duck antibody responses and influenza A virus, several limitations warrant consideration. First, the H3N8 HA used for B-cell isolation was expressed in insect cells and hence lacks the complex glycans typically found on the HA of native influenza virions. As a result, we would fail to isolate some duck antibodies that rely on complex glycans for binding. Second, since our definition of polyreactivity was solely based on the binding activity to SARS-CoV-2 S2 domain and influenza NA, we may have underestimated the total number of polyreactive antibodies. It is possible that more polyreactive antibodies would be identified if a broader panel of antigens were included. Moreover, while our cryo-EM analysis showed that TG0081 utilized a sialylated N-glycan with one LacNAc for binding, it is unclear how frequently duck cells produce this specific glycoform, or if the binding affinity of TG0081 is maintained in the presence of alternative glycoforms. Additionally, some other HA antibodies from ducks (**Table S2**) and even humans^78^ also possess an N-glycosylation site within CDR H3. Whether these antibodies can similarly target the receptor-binding site remains to be determined. Lastly, we acknowledge that the generalization of our findings on H3 subtype to other influenza A subtypes requires additional studies. Overall, this study represents an important step in the molecular understanding of the immune pressure encountered by influenza A virus in their natural reservoir and opens several avenues for future studies.

## ACKNOWLEDGEMENTS

This work was supported by Carl R. Woese Institute for Genomic Biology Postdoctoral Fellowship (H.L.), National Institutes of Health R01 AI165692 (G.-J.B.), Vallee Scholars Program (N.C.W.), Searle Scholars Program (N.C.W.), and Howard Hughes Medical Institute Emerging Pathogens Initiative (J.J.G., B.M.S., and N.C.W.). We thank the Roy J. Carver Biotechnology Center at the University of Illinois Urbana-Champaign for assistance with cell sorting and next-generation sequencing and Frank Vago at the Purdue Cryo-EM Facility for assistance with cryo-EM experiments. We also thank William Lees and Corey Watson for advice on the discovery of immunoglobulin germline genes. We also thank Florian Krammer for providing the H3N2 A/Philippines/2/1982 (X-79) virus, as well as Wilfred van der Donk and Ian Wilson for helpful discussion.

## AUTHOR CONTRIBUTIONS

H.L., W.N.H., B.M.S., R.J.W., and N.C.W. conceived and designed the study. H.L., W.N.H., W.L., D.N., Y.W.H., E.T., T.P., P.S., E.A., A.M., W.J., Q.W.T., A.B.G., E.X.M., D.C.W., M.R.A., A.M., J.J.H., M.T. and G.H. performed the experiments. P.C. fabricated glycan microarrays and performed glycan-binding analysis. G-J.B supervised microarray analysis. B.S., J.J.G., G-J.B, B.M.S., R.J.W., and N.C.W. provided support for the study. H.L. and N.C.W. wrote the paper and all authors reviewed and/or edited the paper.

## METHODS

### Cell culture

HEK293T cells (human embryonic kidney cells, female) and MDCK-SIAT1 cells (Madin-Darby canine kidney cells with stable expression of human 2,6-sialtransferase, female, Sigma-Aldrich) were cultured in Dulbecco’s modified Eagle’s medium (DMEM) with high glucose (Thermo Fisher Scientific) supplemented with 10% heat-inactivated fetal bovine serum (FBS, Thermo Fisher Scientific), 1% penicillin-streptomycin (Thermo Fisher Scientific), and 1× GlutaMax (Thermo Fisher Scientific). Cell passaging was performed every 3 to 4 days using 0.05% Trypsin-ethylenediaminetetraacetic acid (EDTA) solution (Thermo Fisher Scientific). Expi293F cells (human embryonic kidney cells, female, Thermo Fisher Scientific) were maintained in Expi293 Expression Medium (Thermo Fisher Scientific). Sf9 cells (*Spodoptera frugiperda* ovarian cells, female, ATCC) were maintained in Sf-900 II SFM medium (Thermo Fisher Scientific).

### Influenza A virus

Low pathogenic avian influenza A strains A/mallard/Alberta/362/2017 (H3N8), A/mallard/Alberta/328/2014 (H4N6), and A/mallard/Alberta/566/1985 (H6N2) were obtained from routine avian influenza surveillance carried out yearly in Alberta, Canada. Mouse-adapted H3N2 A/Philippines/2/1982 (X-79, 6:2 A/PR/8/34 reassortant) virus was grown in 10-day-old embryonated chicken eggs at 37°C for 48 h and was cooled at 4°C overnight. Cell debris was removed by centrifugation at 4000 ×g for 20 min at 4°C.

### Single-cell sequencing of bulk PBMCs from a mallard duck

Blood samples were collected from a mallard duck (*Anas platyrhynchos*) at Winous Point Shooting Club, Port Clinton, Ohio during the hunting season (October-December) in 2023. The blood samples were collected in tubes containing heparin as an anticoagulant. Blood samples were first centrifuged at 3000 ×g for 10 min at room temperature for plasma collection. The remaining blood was diluted with equal volume of phosphate-buffered saline (PBS), transferred onto the Lymphoprep (STEMCELL Technologies), and centrifuged at 1000 ×g for 20 min to isolate PBMCs, which were subsequently washed with cold RPMI-1640 medium (Thermo Fisher Scientific) three times before storing in cell freezing solution containing 10% dimethyl sulfoxide (DMSO) and 90% FBS at −80°C until used. Single-cell gene expression library was prepared using a Chromium Next GEM Single Cell 5’ Reagent Kits v2 according to manufacturer’s instructions (10x Genomics) and sequenced on an Illumina NovaSeq 6000.

### Single-cell gene expression analysis and cell type identification

Single-cell sequencing data were processed using Seurat^82^. Only cells containing 200–5,000 detected genes were retained for downstream analysis. After log-normalization, 2,000 genes with highly variable expression were identified. Principal component analysis was performed using the top variable genes. The top 30 principal components were used as input for t-distributed stochastic neighbor embedding (t-SNE) analysis. Subsequently, cell clusters were identified using the Seurat function FindNeighbors with default parameters. For each cell cluster, the top differentially expressed genes were matched to the reported cell type markers. Thrombocytes were identified based on the enriched expression of GP9^83,84^, RGS18^85,86^, CCN2^87^, and GREM1^88^. T cells were identified based on the enriched expression of CCR7^89^, CD3E^90^, PLZF^91^, CTLA4^92^. B cells were identified based on the enriched expression of CD79B^93^. NKT cells were identified based on the enriched expression of ADGRG1^94,95^, IL2RB^96,97^, and IL21R^98,99^. Macrophages were identified based on the enriched expression of VCAN and C1QA^100^. Dendritic cells were identified based on the enriched expression of FLT3^101^, PLD4^102^, and JCHAIN^103^. Neutrophiles were identified based on the enriched expression of LTF^104^ and CSF3R^105^.

### Whole-genome sequencing sample preparation and sequencing

Blood samples were collected from a mallard duck (*Anas platyrhynchos*) purchased from Murray McMurry Hatchery (Webster City, IA, USA). PBMCs were extracted as described above. High Molecular Weight (HMW) DNA sample was generated from approximately three million PBMCs using MagAttract HMW DNA Kit (Qiagen). The HMW DNA sample was quantized using Qubit dsDNA Broad Range Assay (Thermo Fisher Scientific) and stored at 4°C to avoid freeze-thaw cycle. Subsequently, the HMW DNA sample was sequenced on a PacBio Revio instrument.

### Genome assembly

PacBio HiFi reads were filtered to remove sequences shorter than 1 kb using seqkit (version 2.6.1)^106^, retaining >95% of reads. Genome size and heterozygosity were estimated from 21-mer frequency distributions generated with KMC (version 3.0.1)^107^ and modeled using GenomeScope2, supporting a diploid genome of ∼1.07 Gb with ∼1% heterozygosity and ∼98× coverage. De novo genome assembly was performed with hifiasm (version 0.19.9)^108^, and both partially phased diploid and haploidized assembly strategies were evaluated. The partially phased diploid assembly produced slightly higher contiguity (N50 ∼22 Mb) but showed reduced BUSCO completeness^109^, consistent with phase-switch errors expected in the absence of auxiliary phasing data. In contrast, a tuned haploidized assembly (−l0 −-hom-cov 90) more effectively purged haplotigs and achieved the highest BUSCO completeness^109^, and was therefore selected as the final assembly. This assembly comprised 483 contigs totaling 1.38 Gb with a contig N50 of 19.9 Mb, without additional scaffolding. Assembly completeness was assessed using BUSCO (version 5.5.0)^109^ with the Avian lineage dataset (aves_odb10), yielding 97.3% complete BUSCOs (93.0% single-copy, 4.3% duplicated), indicating high completeness with limited residual redundancy. Circular contigs observed in the assembly graph are likely organellar sequences, which were not further optimized in this study.

### Annotation of immunoglobulin germline genes

VDJ assemblies from the single-cell VDJ sequencing (see section “Single-cell VDJ sequencing of HA-sorted cells” below) were aligned to the genome using BLASTN (e-value ≤ 10^-5^)^110^ to identify immunoglobulin germline gene sequences. For the heavy chain locus, the *IGHJ* gene was first localized by BLASTN and refined using the flanking recombination signal sequence (RSS) motifs. *IGHD* genes were then defined by targeted identification of paired 12-RSS and 23-RSS motifs located between the *IGHJ* gene and the putative *IGHV* gene region based on the BLASTN result. Subsequently, the *IGHD* and *IGHJ* genes were aligned to each VDJ assembly to identify its *IGHV* nucleotide sequence. The *IGHV* nucleotide sequence of each VDJ assembly was then aligned to the genome to identify *IGHV* genes. Annotation of the light chain locus followed a similar strategy, with the exception that it lacked the D gene. Candidate *IGHV* and *IGLV* genes with less than 200 nucleotides were discarded. The remaining *IGHV* and *IGLV* genes were evaluated by downstream RSS searches with spacer lengths of 22-24 bp, and sequences that translated without stop codons and possessed a valid RSS were classified as functional genes.

### Experimental influenza virus infection of mallard ducks

Male and female mallard ducks (*Anas platyrhynchos*) were purchased from Murray McMurry Hatchery (Webster City, IA, USA) as day-old ducklings and raised to 6-week-old ducklings at the Animal Resource Center at St Jude Children’s Research Hospital. The mallard ducks were leg-banded, and provided feed and water ad libitum. All animal experiments were approved by the Animal Care and Use Committee of St Jude and complied with institutional, National Institutes of Health and Animal Welfare Act policies and regulations. For all experiments, mallard ducks were inoculated by instillation of 10^6^ EID_50_ of virus in a total volume of 500 μL (in PBS) via the natural route (500 μL divided between intranasal, intraocular, intratracheal and oral) on day 0 and day 28. Blood samples were collected in tubes containing heparin as an anticoagulant on day 50 post-infection.

### Expression and purification of recombinant HA proteins

H3 stem, which was previously developed based on A/Finland/486/2004 (H3N2) HA^53^, as well as the HA ectodomain from A/Solomon Island/3/2006 (H1N1), A/California/07/2009 (H1N1), A/Japan/305/1957 (H2N2), A/Hong Kong/1968 (H3N2), A/Singapore/INFIMH-16-0019/2016 (H3N2), A/Darwin/6/2021 (H3N2), A/mallard/Alberta/362/2017 (H3N8), A/cattle/Texas/56283/2024 (H5N1), A/mallard/Alberta/455/2015 (H4N6), A/mallard/Alberta/566/1985 (H6N2), A/duck/Ukraine/1/1963 (H3N8), A/mallard duck/Alberta/635/1983 (H3N5), A/northern pintail/Interior Alaska/9BM8502R0/2009 (H3N1), A/duck/Jiangsu/4/2010 (H3N6), A/mallard/Ohio/14OS0468/2014 (H3N2), and A/Victoria/361/2011 (H3N2) were fused with N-terminal gp67 signal peptide and a C-terminal BirA biotinylation site, thrombin cleavage site, trimerization domain, and a 6x His-tag, and then cloned into a customized baculovirus transfer vector^111^. Subsequently, recombinant bacmid DNA was generated using the Bac-to-Bac system (Thermo Fisher Scientific) according to the manufacturer’s instructions. Baculovirus was generated by transfecting the purified bacmid DNA into adherent Sf9 cells using Cellfectin reagent (Thermo Fisher Scientific) according to the manufacturer’s instructions. The baculovirus was further amplified by passaging in adherent Sf9 cells at a multiplicity of infection (MOI) of 1. Recombinant HA proteins were expressed by infecting 1 L of suspension Sf9 cells at an MOI of 1. On day 3 post-infection, Sf9 cells were pelleted by centrifugation at 4000 ×*g* for 25 min, and soluble recombinant HA proteins were purified from the supernatant by affinity chromatography using Ni Sepharose excel resin (Cytiva) and then size exclusion chromatography using a HiLoad 16/100 Superdex 200 prep grade column (Cytiva) in 20 mM Tris-HCl pH 8.0, 100 mM NaCl. The purified HA proteins were concentrated by Amicon spin filter (Millipore Sigma) and filtered by 0.22 µm centrifuge tube filters (Costar). Concentration of the protein was determined by nanodrop (Fisher Scientific). Proteins were subsequent aliquoted, flash frozen by dry-ice ethanol mixture, and stored at −80°C until used.

HA proteins from the following strains were obtained from BEI Resources: A/New York/55/2004 (H3N2) (cat #: NR-19241), A/Brisbane/10/2007 (H3N2) (cat #: NR-50437), A/Perth/16/2009 (H3N2) (cat #: NR-49734), A/Anhui/1/2013 (H7N9) (cat #: NR-44081) and A/Hong Kong/1073/1999 (H9N2) (cat #: NR-654). HA proteins from the following strains were obtained from Sino Biological: A/pintail duck/Alberta/114/1979 (H8N4), A/duck/Yangzhou/906/2002 (H11N2), A/green-winged teal/ALB/199/1991 (H12N5), A/black-headed gull/Netherlands/1/2000 (H13N8), A/mallard/Astrakhan/263/1982 (H14N5), and A/Australian shelduck/Western Australia/1756/1983 (H15N2).

### Biotinylation of HA proteins

Purified H3 stem and A/mallard/Alberta/362/2017 (H3N8) HA with Y98F mutation were biotinylated using the BirA biotin-protein ligase standard reaction kit according to manufacturer’s instructions (Avidity). Biotinylated H3 stem and H3N8 HA with Y98F mutation were conjugated to streptavidin-BB515 (BD) and streptavidin-PE (Thermo Fisher Scientific). Of note, Y98F mutation was introduced into HA to reduce non-specific HA binding to sialic acid receptors on B cells^112^.

### Single-cell VDJ sequencing of HA-sorted cells

PBMCs were incubated in 1× PBS with 2% FBS, with BB515-conjugated H3 stem and PE-conjugated H3 stem or BB515-conjugated H3N8 HA and PE-conjugated H3N8 HA for 1 h at 4°C. Single BB515^+^PE^+^ B cells were sorted into 5 mL tubes each containing 2 mL of 1× PBS with 2% FBS using a Bigfoot Spectral Cell Sorter. The sorted cells were counted by using 0.4% (w/v) trypan blue stain solution (Thermo Fisher Scientific) under the microscope and loaded on a Chromium Single Cell A Chip (10x Genomics). Subsequently, single-cell lysis and RNA first-strand synthesis were carried out using a Chromium GEM-X Single Cell 5’ Reagent Kit v3 according to manufacturer’s instructions (10x Genomics). The resulting cDNA samples were used for VDJ library construction using a Chromium Single Cell V(D)J Reagent Kit according to manufacturer’s instructions (10x Genomics) with the following primer mixes:

Duck B cell outer primer mix:

- Forward primer: 5’-GAT CTA CAC TCT TTC CCT ACA CGA CGC-3’
- Duck-IgY-outer-R: 5’-TCG CCC AAG GTG GAG CAG ACG GAG CT-3’
- Duck-IgA-outer-R: 5’-CGG TGA GCT GGC TGC TGA CGG TGT A-3’
- Duck-IgM-outer-R: 5’-GGA AGT TGA CAA TGG TGG TGG CCA C-3’
- Duck-IgL-outer-R: 5’-CAG CGT CAG GTA GCT GCT GGC CAT GTA C-3’

Duck B cell inner primer mix:

- Forward primer: 5’-GAT CTA CAC TCT TTC CCT ACA CGA CGC-3’
- Duck-IgY-inner-R: 5’-ACC AGG AGA AGG TGA CGG GA-3’
- Duck-IgA-inner-R: 5’-CGG GGA AGA AGT CGG TGA CGA G-3’
- Duck-IgM-inner-R: 5’-GAG GAG GAG GAG GAA GAG CAG GA-3’
- Duck-IgL-inner-R: 5’-CCG TCA CCG GGC TGG GGT AGA AGT CGC T-3’

The VDJ libraries were then sequenced by an Illumina NovaSeq 6000.

### Analysis of single-cell VDJ sequencing data

Single-cell VDJ sequencing data in FASTQ format were analyzed using Cell Ranger 8.0.0. Briefly, the mallard immunoglobulin germline gene sequences were used as references for both Cell Ranger and IgBlast^113^. After VDJ assembly by Cell Ranger, productive antibody sequences were annotated by IgBlast^113^ Cells with productive heavy and light chains were used for subsequent analyses.

### Phylogenetic analysis of antibody sequences

Chicken germline gene sequences were downloaded from IMGT^49^. Sequence alignment was performed using MAFFT (version 7.526)^114^. Phylogenetic tree was created using FastTree (version 2.1.11)^115^ and visualized using ggtree in R^116^.

### Hydrophobic score of CDR H3

The hydrophobic score for a CDR H3 with a length n was computed as follow:

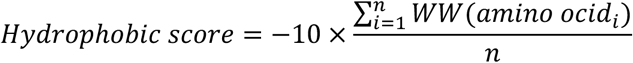

where WW represents the Wimley-White whole residue hydrophobicity scale^117^ and amino acid_i_ represents the amino acid at position i. A higher hydrophobic score represents higher hydrophobicity.

### Expression and purification of IgGs and Fabs

Purified IgGs of the 350 antibody sequences obtained from scVDJ were obtained from Twist Bioscience. For the antibody sequences obtained from phage display antibody screening as well as selected antibodies to be expressed in Fab format, the heavy and light chain genes of the obtained antibody were synthesized as eBlocks (Integrated DNA Technologies), and then cloned into human IgG1 and human kappa or lambda light chain expression vectors using Gibson assembly as described previously^118^. The plasmids of heavy chain and light chain were transiently co-transfected into Expi293F cells at a mass ratio of 2:1 (HC:LC) using ExpiFectamine 293 Reagent (Thermo Fisher Scientific). After transfection, the cell culture supernatant was collected at 6 days post-transfection. The IgGs and Fabs were then purified using a CaptureSelect CH1-XL resin (Thermo Fisher Scientific).

### Enzyme-linked immunosorbent assay (ELISA)

Nunc MaxiSorp ELISA plates (Thermo Fisher Scientific) were utilized and coated with 100 μL of recombinant proteins at a concentration of 1 μg mL^-1^ in a 1× PBS solution. The coating process was performed overnight at 4°C. On the following day, the ELISA plates were washed three times with 1× PBS supplemented with 0.05% Tween 20, and then blocked using 200 μL of 1× PBS with 5% non-fat milk powder for 2 h at room temperature. After the blocking step, 100 μL of IgGs from the supernatant were added to each well and incubated for 2 h at 37°C. The ELISA plates were washed three times to remove any unbound IgGs. Next, the ELISA plates were incubated with horseradish peroxidase (HRP)-conjugated goat anti-human IgG antibody (1:5000, Thermo Fisher Scientific) for 1 h at 37°C. Subsequently, the ELISA plates were washed five times using PBS containing 0.05% Tween 20. Then, 100 μL of 1-Step Ultra TMB-ELISA Substrate Solution (Thermo Fisher Scientific) was added to each well. After 15 min incubation, 50 μL of 2 M H_2_SO_4_ solution was added to each well. The absorbance of each well was measured at a wavelength of 450 nm using a BioTek Synergy HTX Multimode Reader.

### Glycan microarray printing and binding analysis

Glycan microarrays were fabricated and binding analysis was performed as described previously^119,120^. Purified duck antibodies were premixed with AlexaFluor 647-labeled goat anti-human IgG antibody (Fcγ fragment specific, Jackson ImmunoResearch Cat #109-605-008) in TSM buffer (20 mM Tris-HCl pH 7.4, 150 mM NaCl, 2 mM CaCl_2_, 2 mM MgCl_2_, and 1% BSA) with 0.05% Tween-20 in a 2:1 ratio and incubated on ice for 15 min. Precomplexed antibodies were incubated on the microarray slides for 2 h at room temperature. Then, slides were washed by sequential dipping in TSM with 0.05% Tween-20, TSM, and water, followed by centrifugation. The slides were scanned using a GenePix 4000B microarray scanner (Molecular Devices). The images were analyzed using GenePix Pro 7 software (version 7.2.29.2, Molecular Devices).

### Deglycosylation of HA and antibody

Deglycosylation reactions were performed using Protein Deglycosylation Mix II (New England Biolabs). For each reaction, up to 100 μg of purified protein (HA or antibody) was adjusted to a final volume of 40 μL with water, followed by the addition of 5 μL of 10× Deglycosylation Mix Buffer 1 and 5 μL of Protein Deglycosylation Mix II. Reactions were mixed gently and incubated at 25°C for 30 min, then transferred to 37°C and incubated for 16 h.

### Biolayer interferometry binding assay

The binding activity of IgG to H3N8 HA was measured using an Octet Red96e instrument (Sartorius). His-tagged HA ectodomain was loaded onto HIS1K biosensors at 10 μg mL^-1^ in kinetics buffer (1× PBS, pH 7.4, 0.01% w/v BSA and 0.002% v/v Tween 20). The binding experiments were performed with the following steps: 1) baseline in kinetics buffer for 60 s; 2) loading of the HAs for 480 s; 3) baseline for 60 s; 4) association of antibody for 180 s; and 5) dissociation of antibody into kinetics buffer for 180 s. Octet assays were carried out at 25°C. Data were analyzed using the Octet Red Data Analysis software version 9.0. For Fab binding, 1:1 binding model was used to fit the data. Nevertheless, for TG0009 Fab and TG1002 Fab, a 2:1 heterogeneous ligand model was used to improve the fitting due to the weak binding and potential contribution of non-specific binding to the response curve. For IgG binding, a 1:2 bivalent analyte model was used to fit the data.

### Hemagglutination inhibition assays

Hemagglutination inhibition (HAI) assay was performed as described previously^121^. Briefly, 50 µL of H3N2 A/Darwin/6/2021 virus was mixed with 2-fold serial dilutions of plasma samples or antibodies and incubated for 1 h. In some experiments, zanamivir or 1G01 at 100 μg mL^-1^ final concentration was added. After incubation, 50 µL of 1% turkey red blood cells were added to the wells. The highest dilution of the serum that prevented hemagglutination was recorded, and the HAI titer was calculated.

### Virus microneutralization assay

MDCK-SIAT1 cells were seeded in a 96-well, flat-bottom cell culture plate (Thermo Fisher Scientific). The next day, serially diluted monoclonal antibodies were mixed with an equal volume of virus and incubated at 37°C for 1 h. The antibody-virus mixture was then incubated with MDCK-SIAT1 cells at 37°C for 1 h after the cells were washed twice with PBS. Subsequently, the antibody-virus mixture was replaced with Minimum Essential Medium (MEM) supplemented with 25 mM of 4-(2-hydroxyethyl)-1-piperazineethanesulfonic acid (HEPES) and 1 μg mL^-1^ of Tosyl phenylalanyl chloromethyl ketone (TPCK)-trypsin with or without the indicated antibody. The plate was incubated at 37°C for 72 h and the presence of virus was detected by hemagglutination assay.

### Prophylactic protection experiments

Female BALB/c mice at 6 weeks old (n = 5 mice/group) were anesthetized with isoflurane and intranasally infected with 5× median lethal dose (LD_50_) of mouse-adapted H3N2 A/Philippines/2/1982 (X-79, 6:2 A/PR/8/34 reassortant) virus. Mice were given the indicated antibody at a dose of 5 mg kg^-1^ intraperitoneally at 4 h before infection (prophylaxis). Weight loss was monitored daily for 14 days. The humane endpoint was defined as a weight loss of 25% from initial weight on day 0. Of note, while our BALB/c mice were not modified to facilitate interaction with human IgG1, human IgG1 could interact with mouse Fc gamma receptors^121,122^. To determine the lung viral titers on day 3 post-infection, lungs of infected mice were harvested and homogenized in 1 mL of MEM with 1 μg mL^-1^ of TPCK-trypsin with gentleMACS Dissociator (Miltenyi Biotec). Subsequently, viral titers were measured by TCID_50_ (median tissue culture infectious dose) assay. The animal experiments were performed in accordance with protocols approved by UIUC Institutional Animal Care and Use Committee (IACUC).

### Cryo-EM sample preparation and data collection

The purified H3N8 HA ectodomain was mixed with each Fab at a 1:4 molar ratio and incubated at 4°C overnight before purified by size exclusion chromatography. The peak fraction of the Fab-HA complex was eluted in 20 mM Tris-HCl pH 8.0 and 100 mM NaCl, concentrated to around 3 mg mL^-1^, and mixed with n-octyl-β-D-glucoside (Anagrade) at a final concentration of 0.1% w/v for cryo-EM sample preparation. Cryo-EM grids were prepared using a Vitrobot Mark IV machine. An aliquot of 3 μL sample was applied to a 300-mesh Quantifoil R1.2/1.3 Cu grid pre-treated with glow-discharge. Excess liquid was blotted away using filter paper with blotting force 0 and blotting time 3 seconds. The grid was plunge-frozen in liquid ethane. Data collection was carried out on a Titan Krios microscope equipped with Gatan detector. Images were recorded at 81,000× magnification, corresponding to a pixel size of 0.53 Å/pix at the super-resolution mode of the camera. A defocus range of −0.8 μm to −3 μm was used with a total dose of 57.35 e^-^/Å^2^.

### Cryo-EM image processing and model building

Data processing was conducted using CryoSPARC Live (version 4.5)^123^. Movies were subjected to motion correction and contrast transfer function (CTF) estimation, and particles picked with CryoSPARC blob picker followed by 2D classification. For all structures except TG0012, the best classes identified by the blob picker served as templates for CryoSPARC template pickers. The resulting particles underwent multiple rounds of 2D classification to ensure thorough cleanup before proceeding to ab initio reconstruction. The most effective class from the ab initio reconstruction was then subjected to homogeneous refinement, reference-based motion correction, followed by an additional round of homogeneous refinement, local and global CTF estimation, and non-uniform refinement. Finally, all maps except TG0012 were sharpened using DeepEMhancer (version 0.15)^124^.

For model building, ModelAngelo^125^ or AlphaFold^126^ was used to generate an initial model for each Fab, whereas H3N8 HA from PDB 9N4E^127^ was used for HA. In several complexes, particularly those involving Fabs targeting HA glycans, the density for the Fab variable domains was incomplete. In these instances, the density was predominantly well-defined only for the complementarity-determining regions (CDRs) directly interacting with the glycans. To account for this, AlphaFold^126^ models of the Fabs were generated and docked into the maps. These models exhibited high conformational agreement with the available density. The full AlphaFold structures were retained in the final models despite the partial definition of the rest of the variable domain. To improve the resolution of these interfaces, local refinement was performed using masks focused on the epitope-paratope boundary to enhance the visualization of specific sidechain and glycan interactions. These models were fitted into the cryo-EM density map using UCSF Chimera^128^. They were then subjected to multiple rounds of manual refinement in Coot (version 0.9.8)^129^ and real-space refinement in Phenix (version 1.21.1-5286)^130^. The iSOLDE tool within UCSF ChimeraX was also used^131^. This process was iterated for several cycles until no significant improvement of the model was observed. The buried surface area (BSA) of the epitope and paratope was computed using PDBePISA^132^.

### HA sequence analysis

A total of 2422 full-length duck H3 HA sequences and 140,296 full-length human H3 HA sequences were downloaded from Global Initiative for Sharing Avian Influenza Data (GISAID)^133^.

To avoid temporal sampling bias, we sampled at most 10 sequences per year, which resulted in 364 duck H3 HA sequences and 527 human H3 HA sequences. Sequence alignment was performed using MAFFT (version 7.526)^114^. Sequences with ambiguous amino acids were discarded. Sequence logos were generated by Logomaker in Python^134^. Phylogenetic tree was created using FastTree (version 2.1.11)^115^ and visualized using ggtree in R^116^.

## CODE AVAILABILITY

Computer scripts for annotating the mallard immunoglobulin germline genes have been deposited to https://github.com/nicwulab/mallard_VDJ_analysis.

## DATA AVAILABILITY

Raw sequencing data and the haploid assembly of the mallard genome have been deposited in the NIH Short Read Archive under accession number: PRJNA1394649 and PRJNA1393375, respectively. Cryo-EM maps have been deposited in the Electron Microscopy Data Bank with accession numbers: EMD-75031, EMD-75032, EMD-75033, EMD-75197, EMD-75199, EMD-75034, EMD-75200, EMD-75201, EMD-75202, EMD-75203, EMD-75204, EMD-75205, EMD-75206, EMD-75207, EMD-75208, EMD-75209, EMD-75210, and EMD-75717. The refined models have been deposited in the Protein Data Bank with accession numbers: 10AO, 10AQ, 10AR, 10IL, 10IN, 10AS, 10IO, 10IP, 10IQ, 10IR, 10IS, 10IT, 10IU, 10IV, 10IW, 10IX, 10IY, and 11IL.

