## Supplementary material for "Structural insights into antibody responses against influenza A virus in its natural reservoir": Figure S1-S12

A

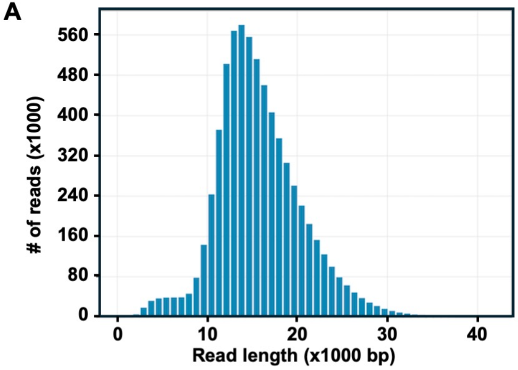


B

| Sequencing statistics | |
|---|---|
| Total number of bases (bp) | 105,725,782,409 |
| Total number of reads | 6,723,167 |
| Max read length (bp) | 55,452 |
| Min read length (bp) | 69 |
| Mean read length (bp) | 15,726 |
| **Post-assembly analysis** | |
| Number of contigs | 481 |
| N50 (bp) | 19,862,632 |
| BUSCO completeness | 97.3% |
| BUSCO duplication | 4.3% |

C

```
                                               CDR H1                    CDR H2
IGHV1-1   AATLDESGGGLVSPGGSLTLVCKGSGYTFSS-YGMGWVRQAPGKGLEYVASINSSGSSTYYAPAVKGRFTISRNNGQSTLTLQMNSLKAEDTATYYCAK
pseudogenes
IGHV1-3   AATLDKTGGGLVSPGGSLTLVCKGSDFSFKD-YTMEWLRQAPGKGLEYVAGISYSSSYTVYAPAVQGRFTISRDNAQSTVTLQMNSLKAEDTGTYYCAK
IGHV1-4   AMTLDESGGGRVSPGGSLTLVCKGSGFTFSS-YTMQWVRQAPRKGLEWVARISNTGSTTNYAPAVKGRFTISKNDGQSTLTLQMNSLKAEDTATYYCAK
IGHV1-5   ALMLDESGGGLVSPGGSLTLVCKGSGFTFSY-YDMFWVRQAPGKALEFVAGITSSGSYTVYAPAVKGRFTISKNDGQSTATLQMNSLKAEDTATYYCAK
IGHV1-8   AETLDESGGGLVSPGGSLTLVCKGSGFTFSS-YYLDWVRQAPGKELEWVATIKYNGRTGWYLPEVEGRFTISRNNGQSTVTLQMNSLKAEDTATYYCTR
IGHV1-9   AETLDESGGGLVSPGGSLTLVCKASGFDFSS-NDMGWMRQAPGKGLEVVAVINNDGGSTWYAAAVKGRFTISRDNGQTVFFLLMSDLKAEDTATYYCVK
IGHV1-11  AATLDESGGTLMNPGGSLTLVCKGSGYTFSD-YRIYWVRQAPGKGLEFFAGIESDGD-TFYAPAVKGRFTVSRNNGQSTATLQMNSLKAEDTATYYCA-
IGHV1-13  ALMLDESGGTLVSPGGSLTLVCKGSDFTFNY-YGMSWVRQAPGKGLELVAGIDDDGSYTEYAPAVKGRFTISRNNGQSTATLQMNSLKAEDTATYYCAK
IGHV1-16  AATLDESGGGLVSPGGSLTLVCKGSGFTFSS-WGMQWVRQAPGKGLEWVAQISSTGSATYYAPAVKGRFTISRNNGQSTLTLQMNSLKAEDTATYYCAK
IGHV1-18  AATLDESGGGLVSPGGSLTLVCKGSGFDFSS-TDMGWMRQAPGKGLEYVASIYSSSSYTDYASVVQGRFTISRNNGQSTLTLQMNSLKAEDTATYYCTK
IGHV1-19  AATLDESGGGLVSPGGSLTLVCKGSGFTFSS-YNMQWVRQAPGKGLEYVAEISSSGGSTAYAPAVKGRFTISRNNGQSTATLQMNSLKAEDTATYYCVR
IGHV1-21  AETLDESGGGRVSPGGSLTLVCKGSGYTFSD-YGMRWVRQAPGKGLEDIAAISKDGSWTGYASAVKGRFTISRNNGQSTATLQMNSLKAEDTATYYCAK
IGHV1-23  AETLDESGGGLVSPGGSLTLVCKASGFTFSS-YNMEWVRQAPGKGFEFVAQINPGGSYTGYASAVKGRFTISRNNGQSTVTLQMNSLKAEDTATYYCAR
IGHV1-26  AATLDESGGGLVSPGGSLTLVCKASGFTFSSNY-MYWVRQAPRKGLEFVAYISSSGSYTNYAPAVKGRFTISRNNGQSTATLQMNSLKAEDTATYYCAR
IGHV1-27  AATLDESGGGLVSPGGSLTLVCKGSGFTFSS-NYMSWVRQAPGKGLEFVAYISTGGSSTYYAPAVKGRFTISKNDGQSTATLQMNSLKAEDTATYYCAK
IGHV1-29  ALMLDESGGGLVSPGGSLTLVCKGSDFTFSY-YGMGWMR*APGKGLEYVAGITDSGGYTEYASAVKGRFTISRNNGQSTATLQMNSLKAEDTATYYCAK
IGHV1-30  AETLDESGGGLVSPGGSLTLVCKGSGYTFSS-YGMYWVRQAPGKGLEYVGSIHSSGSGTDYASAVQGRFTISRNNGQSTATLQMNSLKAEDTATYYCAK
IGHV1-31  AETLDESGGGRVSPGGSLTLVCKGSGYTYSE-YRIYWVRQAPGKGLEYVAQIGYSGSSTHYAPAVEGRFTISRNDGQSTATLQMNSLKAEDTATYYCAR
IGHV1-33  AMTLDESRGGLVSPGGSLTLVCKGSGFDFSS-NDMEWVRQAPRKGLEFVATIWSD-GNTWYLPAVKGRFTISRDNGQSLLFLLMNDLKAEDTATYYCAR
IGHV1-34  ALMLDESGGGLVSPGGSLTLVCKGSGFTFGN-YSMFWVRQALGKGLEYVAAIN-TGSTTNYAPAVKGRFTISRNNGQSTATLQMNSLKAEDTATYYCTK
IGHV1-37  AETLDESGGGLVSPGGSLTLVCKGSGFTFSS-YAMYWVRQAPGKGLEYVAAINTDGT-TYYASAVQGRFTISRSYWYSTATLQMNSLKAEDTATYYCAK
IGHV1-38  AETLDESGGGRVSPGGSLTLVCKGSGYTFSD-YGMRWVRQAPGKGLEDIAAISKDGSSTTYAPAVKGRFTISKNDGQSTVTLQMNSLKAEDTATYYCAK
IGHV1-40  AANLDETGGGLVKPGGSLTLVCKGSGYTFSSNYNTEWVRQAPRKGLEFVAGISGSGSYTGYAPAVEGRFTISRNNGQSTLTLQMNSLKAEDTATYYCAK
IGHV1-49  AATLDESGGGRVSPGGSLTLVCKASGFTFSS-YGMSWVRQAPGKGLEYVAGITGSGGYTDYAPAVKGRFTISRNDGQSTLTLQMNSLKAEDTATYYCAR
IGHV1-50  AMTFDESGGGLVSPGGSLTLVCKGSGYTFSS-YYLDWVRQAPGKELEWVATIKYNGRTGWYLPEVEGRFTISRNDGQSTATLQMNSLKAEDSATYYCTR
IGHV1-51  AETLDESGGGLVSPGGSLTLVCKGSGFTFSS-YGMNWVRQAPGKALEFVAGIDNSGSYTEYAPAVQGRFTISRNNGQSTATLQMNSLKAKDTGTYYCAK
IGHV1-53  AETLDESGGTLVNPGGSLTLVCKASGFTFSS-YYLDWVRQAPGKELEWVATIKYNGRTGWYLPEVEGRFTISRNDGQSTLTLQMNSLKAEDTATYYCTR
IGHV1-55  AATLDESGGGLVSPGGSLTLVCKGSGFTFSS-YSMGWIRQAPGKGLEWVATIYSSGSSPWYASAVKGRFTISRNNGQSTVTLQMNSLKAEDTATYYCAK
IGHV1-56  AETLDESGGGLVSPGGSLTLVCKGSGYTFSSNYNTEWVRQAPRKGLEFVAYISSSGGYTSYAPAVKGRFTISRNDGQSTATLQMNSLKAEDTATYYCAR
IGHV1-58  AATLDESGGGLVSPGGSLTLVCKGSGFTFST-YNMAWMRQAPGKGLEYVAAIN-TGGTTYYAPAVKGRFTISRNNGQSTATLQMNSLKAEDTATYYCAK
          27 31 32 32a 33 34 35 50 51 52 52a 53 54 55 56 57 58 59 60 61 62 63 64 65 71
```

D

```
                                            CDR H1                    CDR H2
Mallard IGHV1-1   AATLDESGGGLVSPGGSLTLVCKGSGYTFSSYGMGWVRQAPGKGLEYVASINSSGSSTYYAPAVKG
Chicken IGHV1-1   AVTLDESGGGLQTPRGALSLVCKASGFTFSSYGMGWVRQAPGKGLEWVAGIGSSGSGTAYGSAVKG

Mallard IGHV1-1   RFTISRNNGQSTLTLQMNSLKAEDTATYYCAK
Chicken IGHV1-1   RATISRDNGQSTVRLQLNNLRAEDTGTYYCAK
```

E

```
                                         CDR L1              CDR L2
Mallard IGLV1-1   QAALTQPASKSVNPGDTVQITCSGGGSYYGWFQQKTPGTGPVTVIYDNTNRPSGIPSRFSASTSGS
Chicken IGLV1-1   QAALTQPSSVSANPGETVKITCSGDRSYYGWYQQKAPGSAPVTVIYANTNRPSDIPSRFSGSKSGS

Mallard IGLV1-1   VSTLTITGVQAEDEAVYYCG
Chicken IGLV1-1   TATLTITGVQADDEAVYYCG
```

**Figure S1. Whole-genome sequencing and identification of immunoglobulin germline gene.** **(A)** The length distribution of PacBio HiFi reads. **(B)** Summary statistics of the sequencing run and the subsequent genome assembly. **(C)** Sequence alignment of representative mallard *IGHV* genes. Positions of interest are indicated at the bottom. **(D)** Sequence alignment of functional *IGHV1-1* genes from mallard and chicken. **(C-D)** Red: differences from mallard *IGHV1-1*. **(E)** Sequence alignment of functional *IGLV1-1* genes from mallard and chicken. Red: differences from mallard *IGLV1-1*.

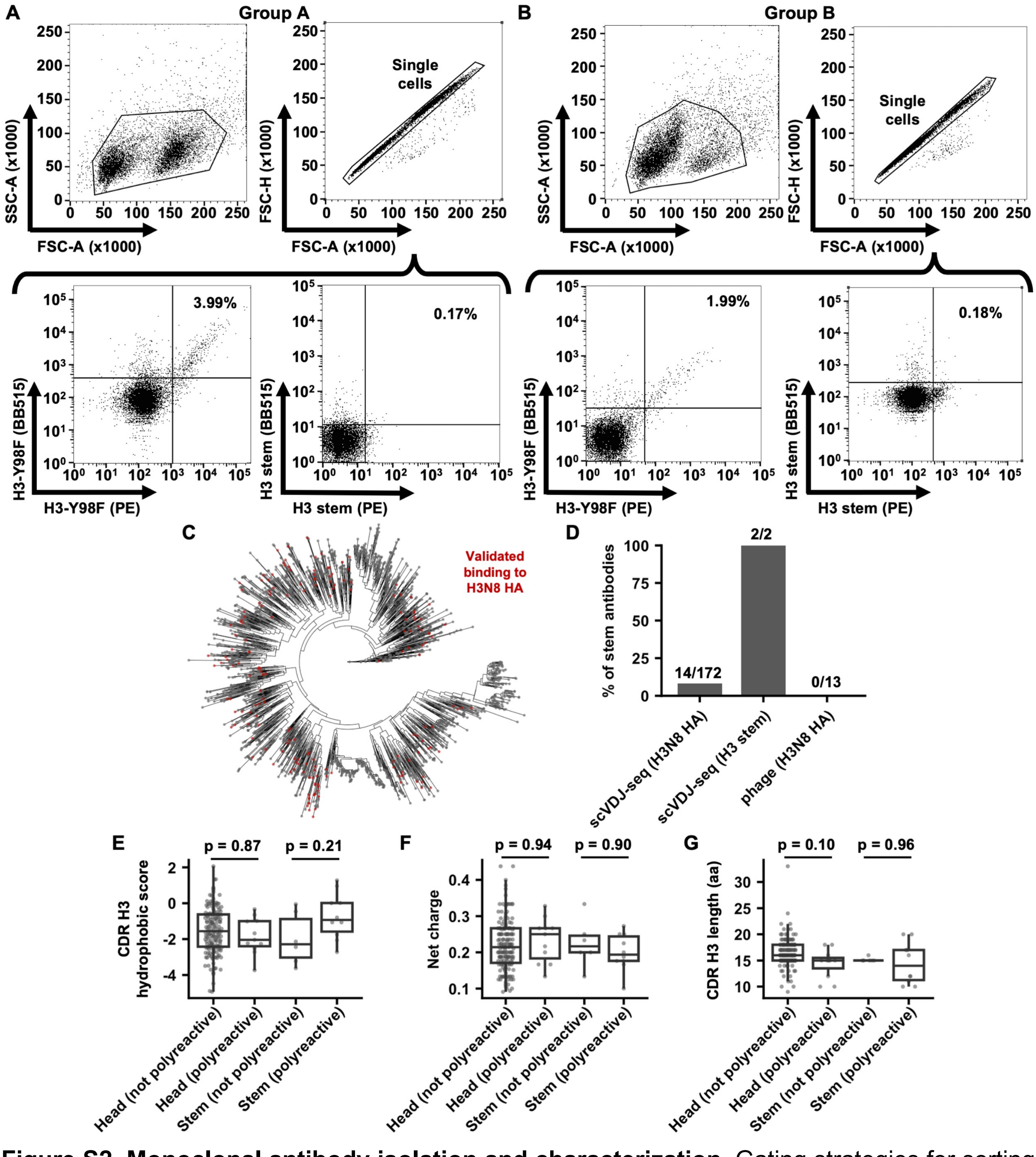


**Figure S2. Monoclonal antibody isolation and characterization.** Gating strategies for sorting H3N8 HA-specific and H3 stem-specific cells from **(A)** group A mallards and **(B)** group B mallards. **(C)** Phylogenetic tree of 2417 duck monoclonal antibody sequences. Red: antibodies with validated binding activity to H3N8 HA. **(D)** Frequency of stem-binding antibodies among those isolated via different methods. **(E)** Hydrophobicity score, **(F)** net charge, and **(G)** CDR H3 length in the indicated categories of antibodies. The p-values were computed by two-tailed Student's t-tests. For the boxplot, the middle horizontal line, lower, and upper hinges represent the median, the first and third quartiles, respectively. The upper and lower whiskers extend to the highest data point within 1.5× inter-quartile range (IQR) of the third quartile, and the lowest data point within 1.5× IQR of the first quartile, respectively. Each data point represents one antibody.

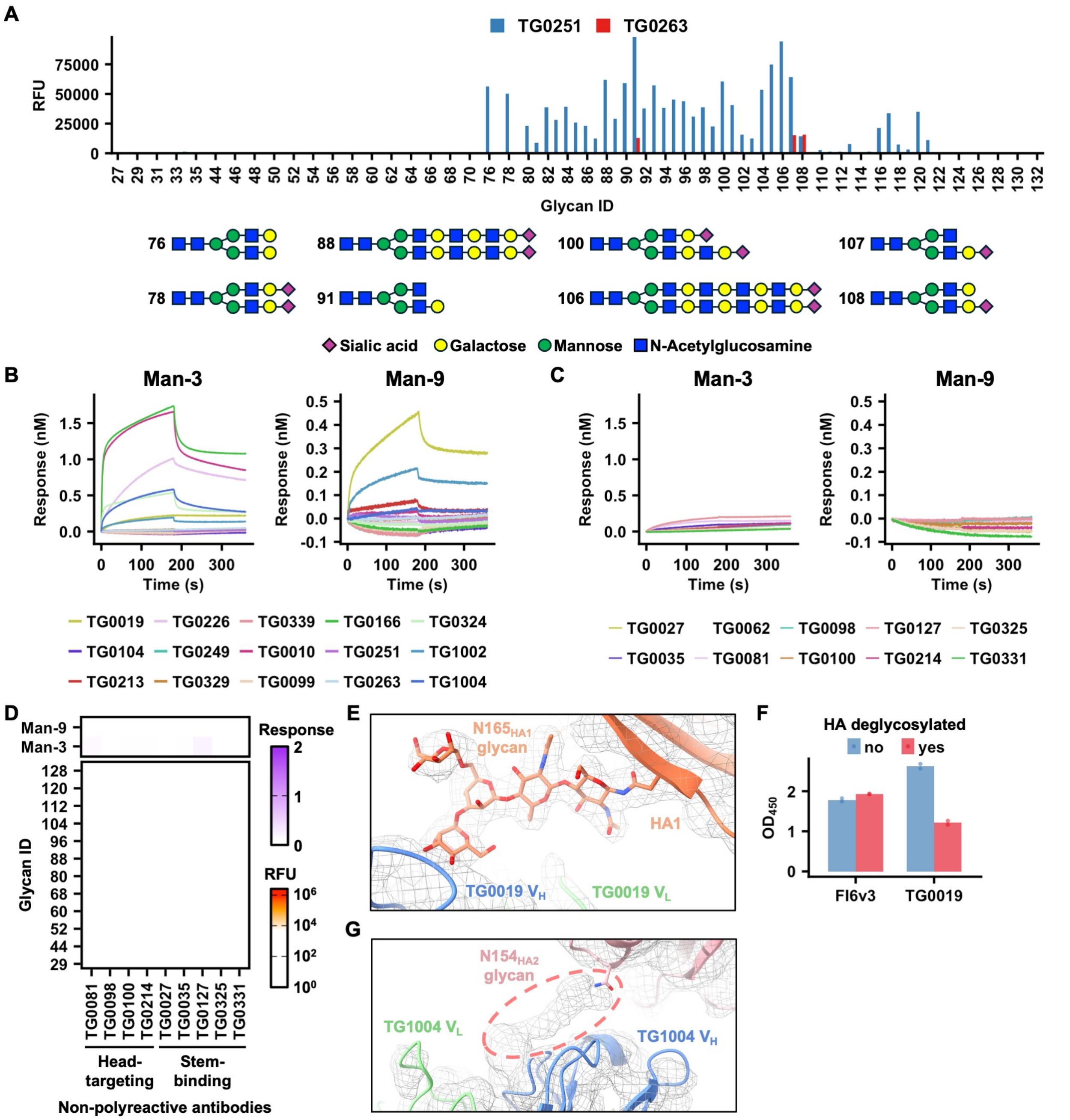


**Figure S3. Glycan-binding activity of polyreactive duck antibodies. (A)** Glycan array data for TG0251 (blue) and TG0263 (red). Glycan diagrams for representative glycans are shown at the bottom according to the Symbol Nomenclature for Glycans recommended by the NLM[1,2]. **(B-C)** Binding activity of the indicated **(B)** polyreactive antibodies and **(C)** non-polyreactive antibodies to mannose-3 (Man-3) and mannose-9 (Man-9) was measured by biolayer interferometry. **(D)** Glycan-binding activity of the indicated non-polyreactive antibodies was tested by glycan microarray (bottom panel, red) and biolayer interferometry (upper panel, purple). Man-9: mannose-9. Man-3: mannose-3. **(E)** Cryo-EM electron density map (grey mesh) of H3N8 HA in complex with TG0019. **(F)** The binding activity of FI6v3 and TG0019 to H3N8 HA with (red) or without (blue) deglycosylation was measured by ELISA. **(G)** Cryo-EM electron density map of H3N8 HA in complex with TG1004 is shown as a grey mesh. The HA trimer (PDB 9N4E)[3] and an AlphaFold[4] model of the TG1004 were docked into the density using ChimeraX[5]. Unassigned density near $N154_{HA2}$, potentially representing an N-linked glycan, is outlined with pink dashes. Due to the map resolution (4.16 Å), glycans could not be modeled with high confidence.

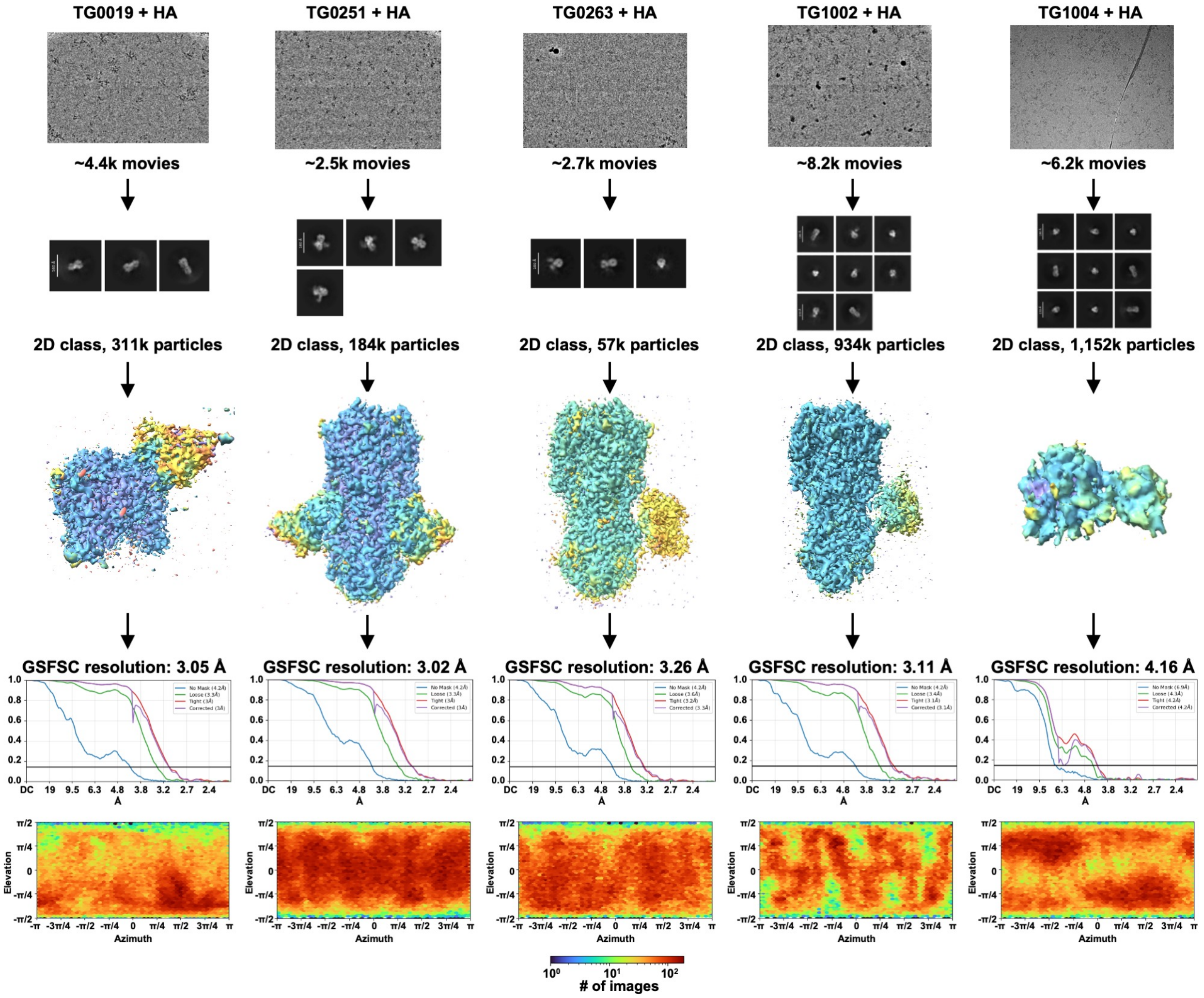


**Figure S4. Cryo-EM data processing for the structures of TG0019, TG0251, TG0263, TG1002, and TG1004 Fabs in complex with H3N8 HA.** Representative micrographs, representative 2D class averages, local resolution maps, Fourier shell correlation (FSC) curves, and Azimuth plots are shown.

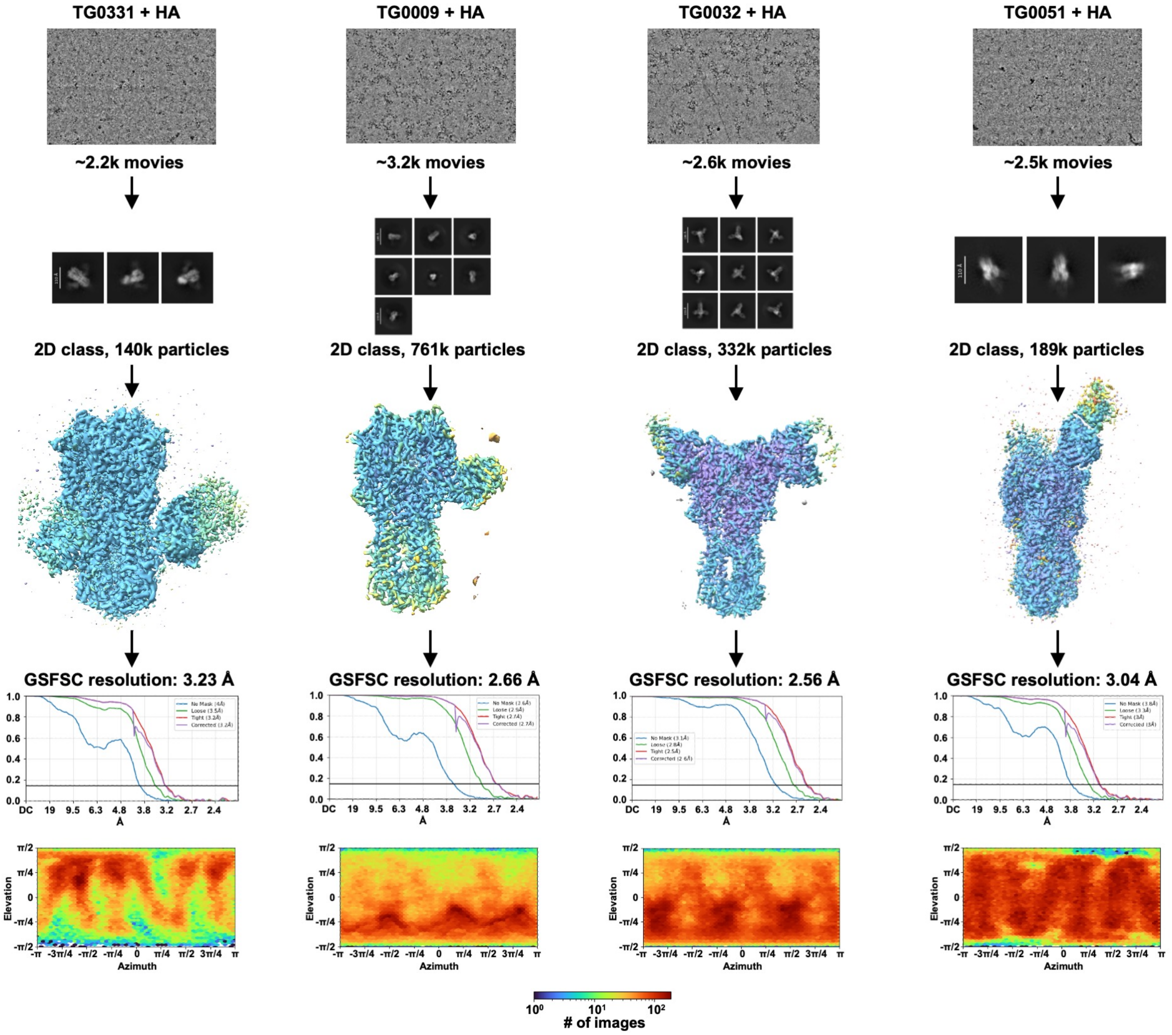


**Figure S5. Cryo-EM data processing for the structures of TG0331, TG0009, TG0032, and TG0051 Fabs in complex with H3N8 HA.** Representative micrographs, representative 2D class averages, local resolution maps, Fourier shell correlation (FSC) curves, and Azimuth plots are shown.

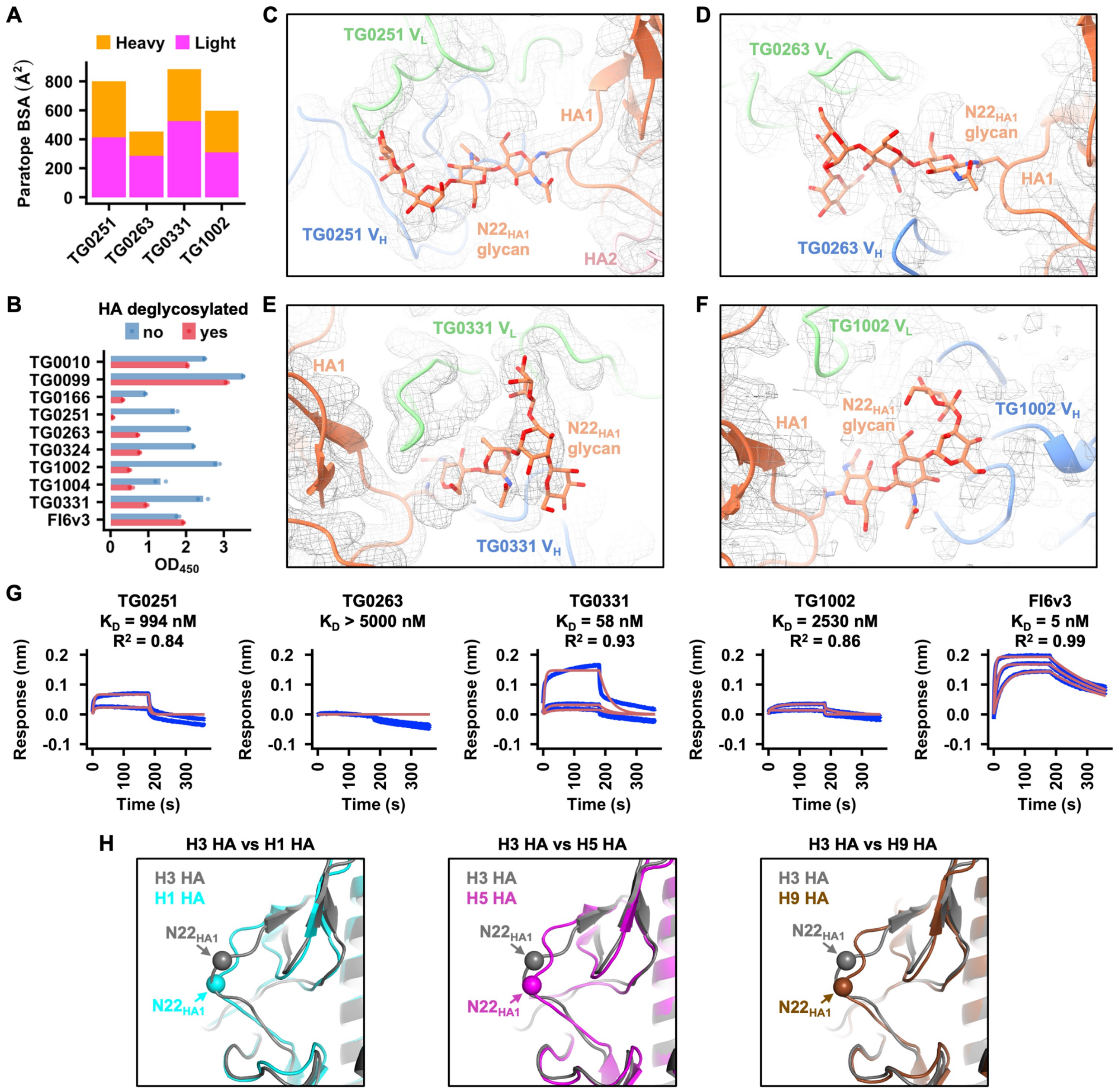


**Figure S6. Structural characterization and binding activity of stem-binding duck antibodies. (A)** Paratope composition of the indicated antibodies analyzed by buried surface area (BSA) contributions of the heavy and light chains to HA interaction. **(B)** Binding activity of the indicated antibodies to H3N8 HA with (red) or without (no) deglycosylation was measured by ELISA. FI6v3 is a known stem-binding human antibody that does not rely on glycans for HA binding[6]. **(C-F)** Cryo-EM density maps (grey mesh) of H3N8 HA in complex with **(C)** TG0251, **(D)** TG0263, **(E)** TG0331, and **(F)** TG1002 are shown. **(G)** Binding activity of the indicated antibodies in Fab format to H3N8 HA was measured by biolayer interferometry. Y-axis represents the response. Blue and red lines represent the response curve and the 1:1 binding model, respectively. Binding kinetics were measured for 300 nM, 100 nM, and 33 nM of each Fab. (H) Structural alignment between H3 HA and three different group 1 HAs, namely H1 HA (PDB 6CF7)[7], H5 HA (PDB 9DIQ)[8], and H9 HA (PDB 1JSH)[9]. The Cα atoms of the indicate residues is shown as spheres.

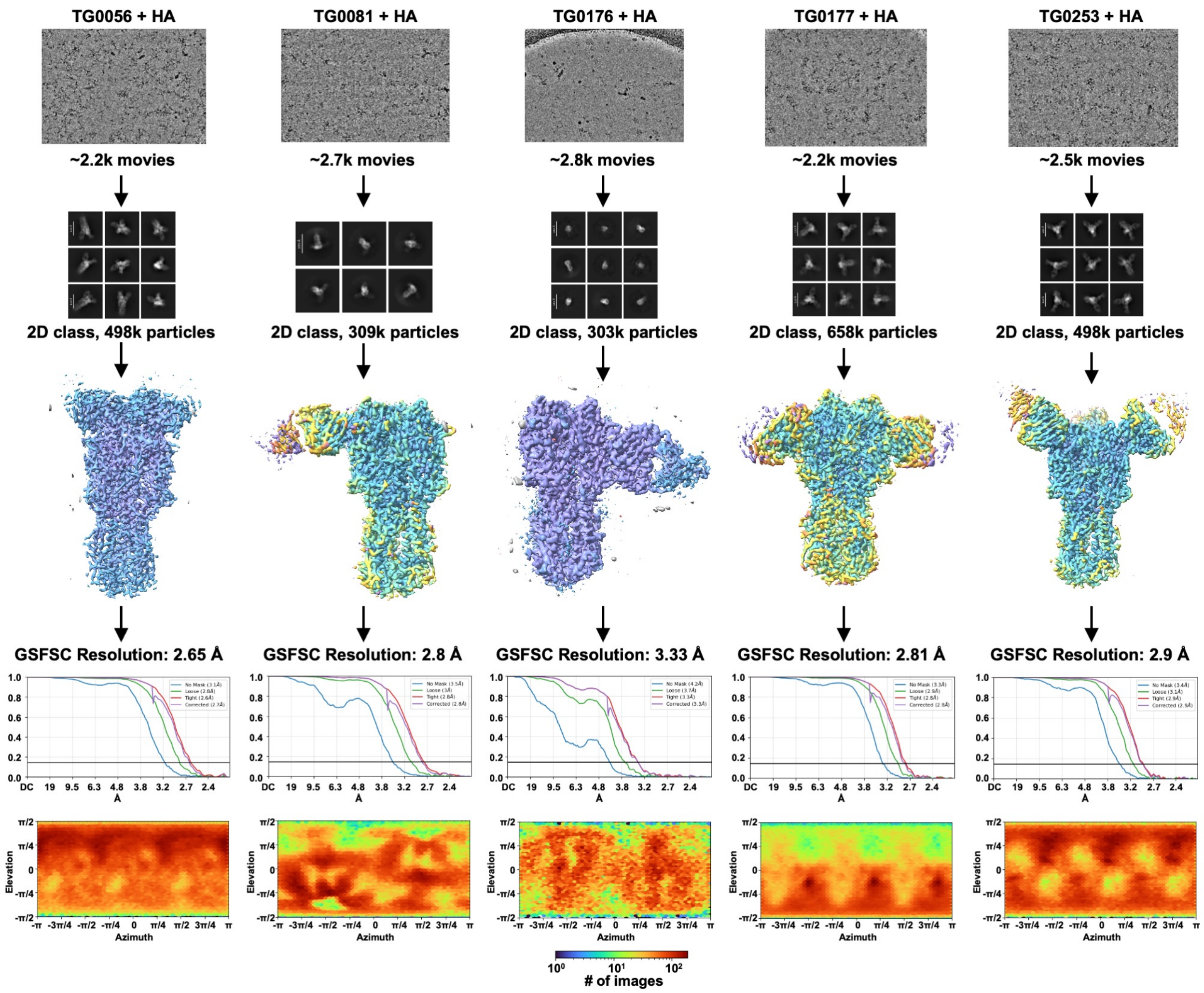


**Figure S7. Cryo-EM data processing for the structures of TG0056, TG0081, TG0176, TG0177, and TG0253 Fabs in complex with H3N8 HA.** Representative micrographs, representative 2D class averages, local resolution maps, Fourier shell correlation (FSC) curves, and Azimuth plots are shown.

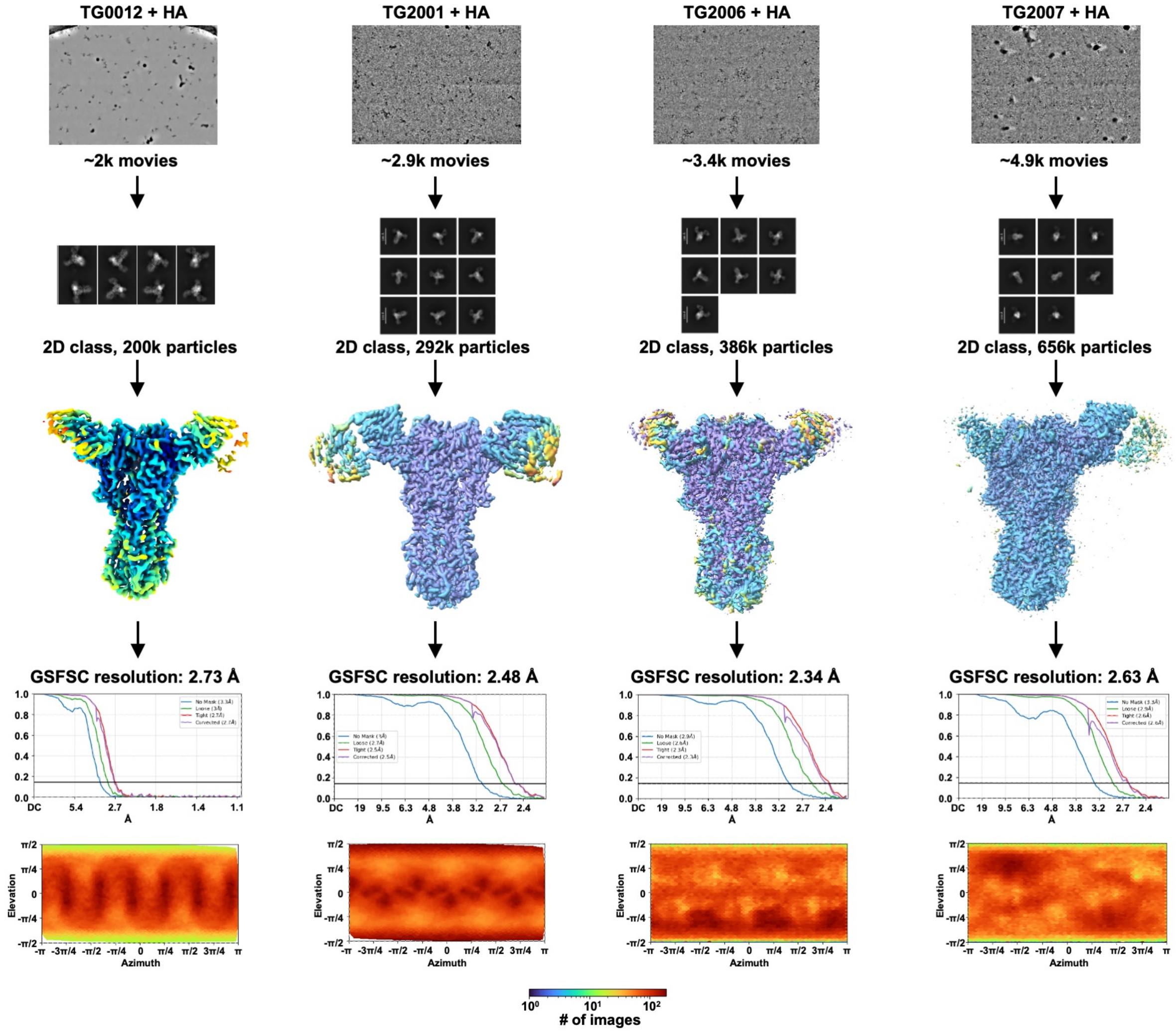


**Figure S8. Cryo-EM data processing for the structures of TG0012, TG2001, TG2006, and TG2007 Fabs in complex with H3N8 HA.** Representative micrographs, representative 2D class averages, local resolution maps, Fourier shell correlation (FSC) curves, and Azimuth plots are shown.

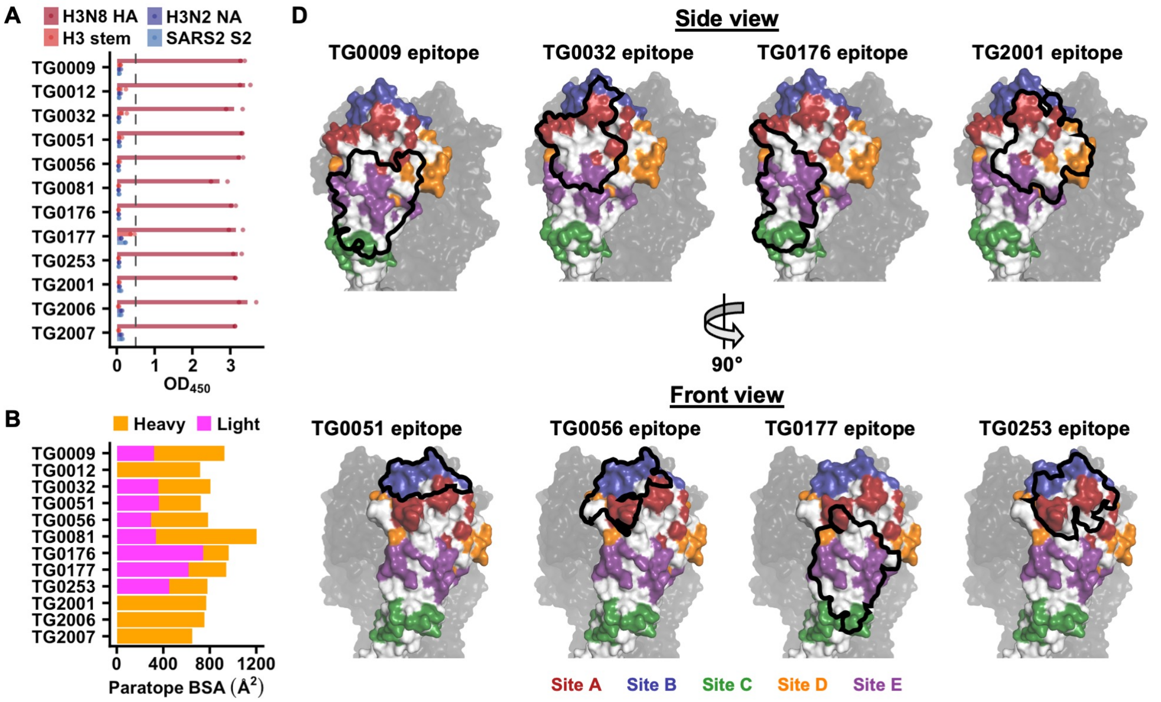


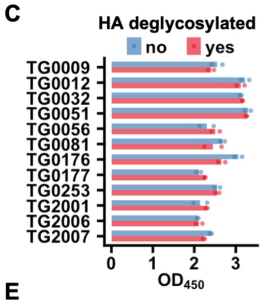


**E**

| Antibody | $k_{on}$ ($\mu M^{-1}\ s^{-1}$) | $k_{off}$ (1/s) | $K_D$ (nM) |
|---|---|---|---|
| TG0009 | 0.07 | 0.114 | 1650 |
| TG0032 | 0.87 | 0.006 | 7 |
| TG0051 | 0.07 | 0.013 | 172 |
| TG0056 | 0.12 | 0.001 | 12 |
| TG0081 | 1.04 | 0.051 | 49 |
| TG0177 | 0.06 | 0.046 | 772 |
| TG2001 | 1.02 | 0.009 | 9 |
| TG2006 | 0.17 | 0.011 | 66 |
| TG2007 | 0.31 | 0.018 | 60 |

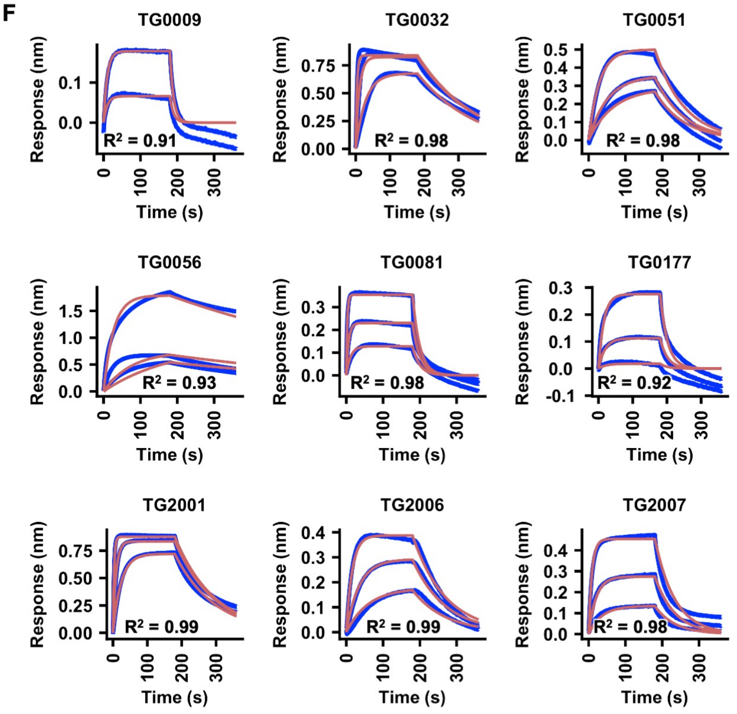


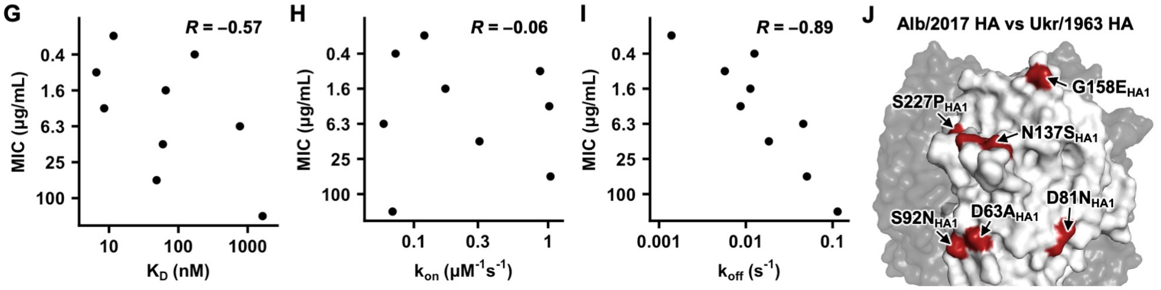

**Figure S9. Structural characterization and binding activity of head-targeting duck antibodies. (A)** Binding activity of the indicated antibodies against the indicated antigens was measured by ELISA. Each bar represents the average of two replicates. Each data point represents one replicate. H3N8 HA: A/mallard/Alberta/362/2017 (H3N8) HA. H3 stem: a stabilized HA stem construct previously developed based on A/Finland/486/2004 (H3N2) HA[10]. H3N2 NA: A/Moscow/10/1999 (H3N2) NA. SARS2 S2: SARS-CoV-2 S2 domain. **(B)** Paratope composition of indicated antibodies. **(C)** The binding activity of the indicated antibodies to H3N8 HA with (red) or without (blue) deglycosylation was measured by ELISA. **(D)** Epitopes of the indicated antibodies are outlined on one HA protomer with the major antigenic sites indicated. The other two HA protomers are shown in black. **(E)** The $K_D$, on-rate ($k_{on}$), and off-rate ($k_{off}$) for the indicated antibodies are shown. **(F)** Binding activity of the indicated antibodies in Fab format to H3N8 HA was measured by biolayer interferometry. Y-axis represents the response. Blue lines represent the response curve. Red lines represent the 1:1 binding model, except for the TG0009 data, where a 2:1 heterogeneous ligand model was applied due to potential non-specific binding (**see Methods**). Binding kinetics were measured for 300 nM, 100 nM, and 33 nM of each Fab. **(G-I)** The relationship between neutralization activity as determined by microneutralization assay and binding kinetics, including **(G)** $K_D$, **(H)** on-rate ($k_{on}$), and **(I)** off-rate ($k_{off}$) was analyzed for nine head-targeting duck antibodies. Binding kinetics were measured using Fabs. MIC: minimum inhibitory concentration. **(J)** The amino acid mutations in the head domain of A/duck/Ukraine/1/1963 (H3N8) HA, relative to A/mallard/Alberta/362/2017 (H3N8) HA, are highlighted on the structure.

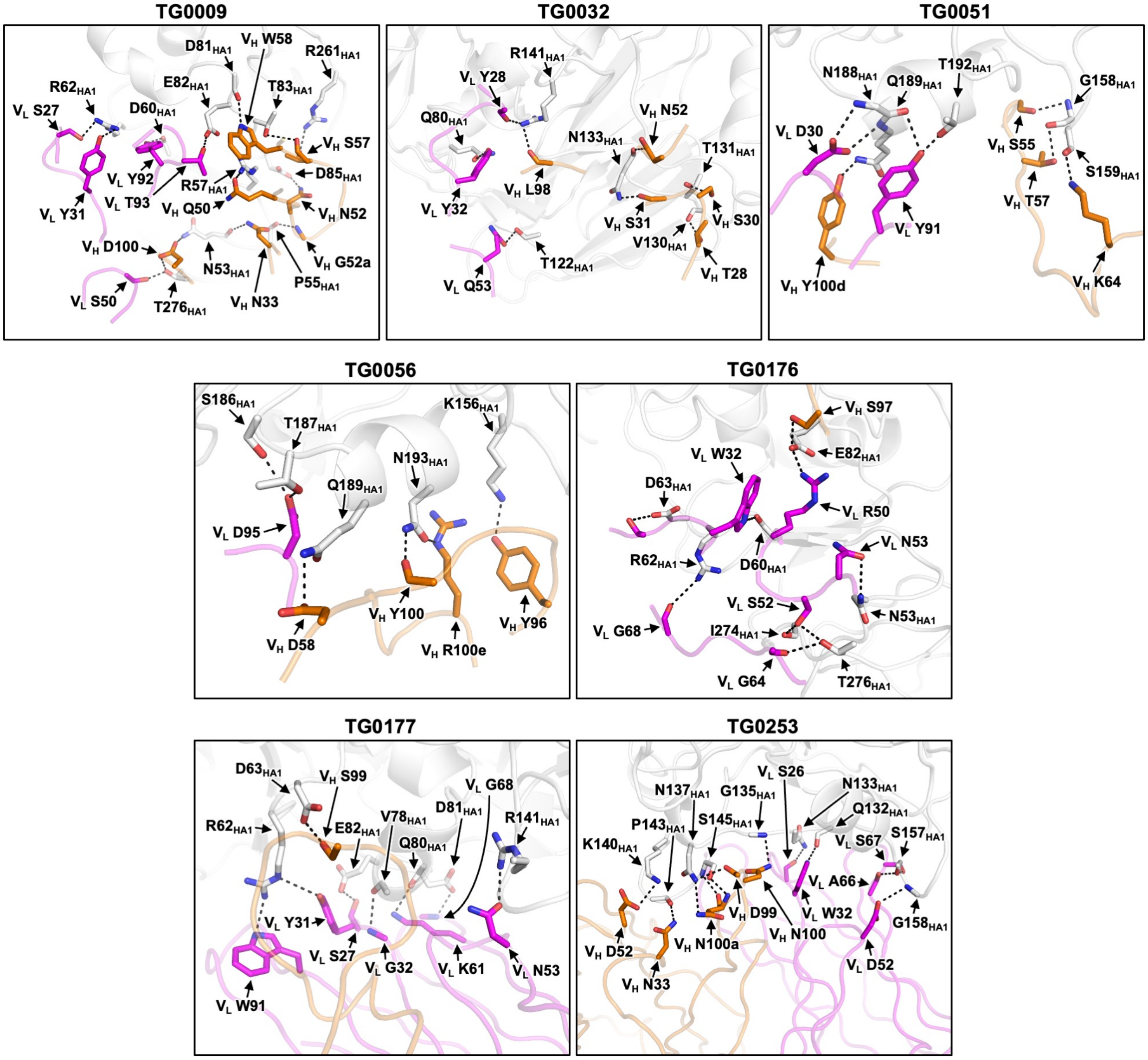


**Figure S10. Interaction between representative head-targeting duck antibodies and H3N8 HA.** Key residues at the interfaces between the indicated antibodies and HA are shown. White: HA. Orange: antibody heavy chain. Magenta: antibody light chain. Black dashed lines represent H-bonds or electrostatic interactions.

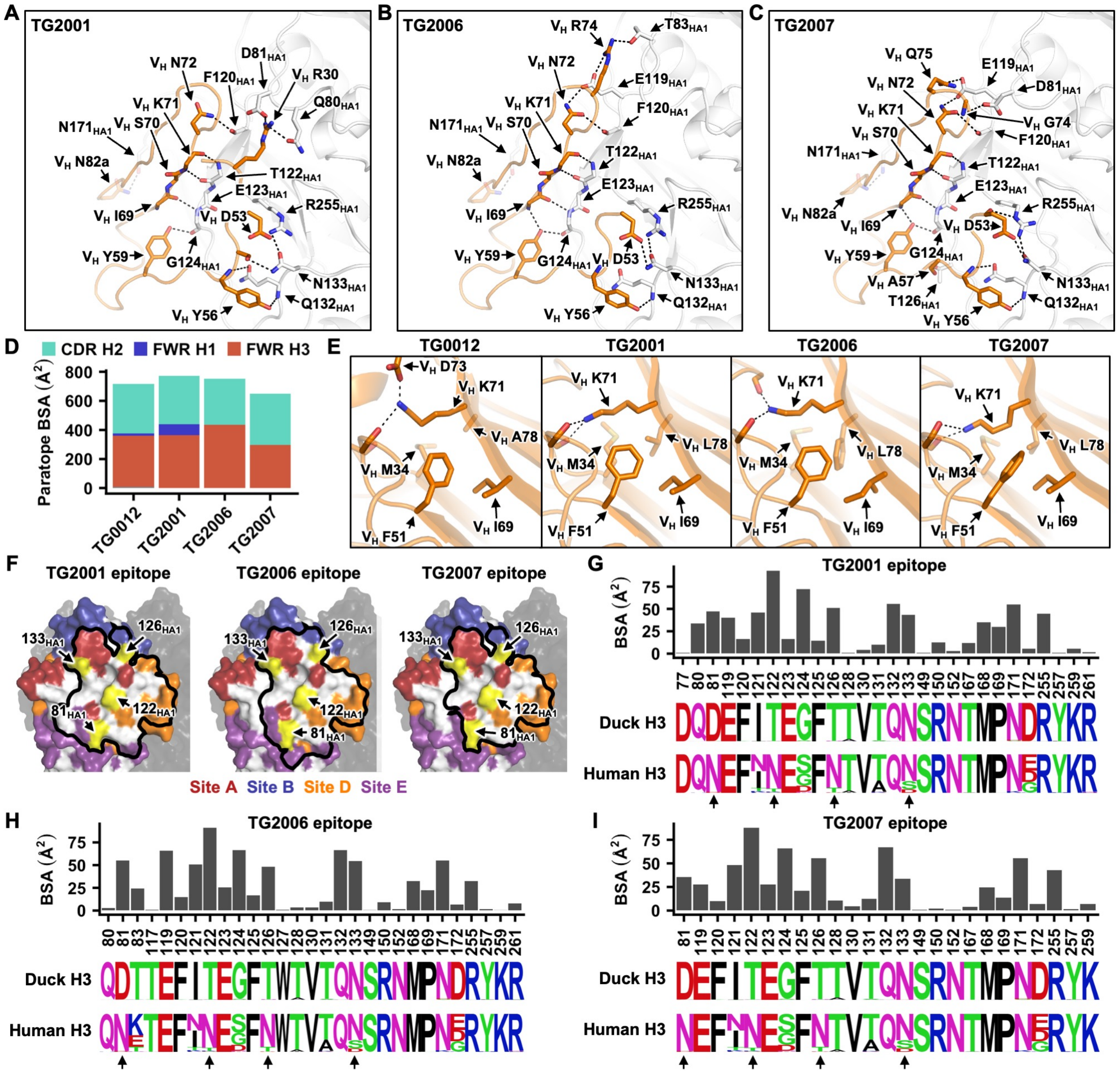


**Figure S11. Structural characterization of a convergent antibody binding mode. (A-C)** Key residues at the interfaces **(A)** between TG2001 and HA, **(B)** between TG2006 and HA, and **(C)** between TG2007 and HA are shown. White: HA. Orange: antibody heavy chain. Black dashed lines represent H-bonds or electrostatic interactions. **(D)** Paratope composition of indicated antibodies. **(E)** Intramolecular interactions involving the side chains of $V_H$ K71 in TG0012, TG2001, TG2006, and TG2007 are shown. Black dashed lines represent H-bonds. **(F)** Epitopes of TG2001, TG2006, and TG2007 are outlined on one HA protomer with the major antigenic sites indicated. The other two HA protomers are shown in black. N-glycosylation sites with high occurrence frequency in human H3 strains are shown in yellow. **(G-I)** The bar charts indicate the buried surface area (BSA) of each residue in the epitopes of **(G)** TG2001, **(H)** TG2006, and **(I)** TG2007 upon binding. The sequence logos represent the sequence diversity of each epitope residue in duck H3 strains from 1963 to 2025, as well as human H3 strains from 1968 to 2025. N-glycosylation sites present in human H3 strains but not duck H3 strains are indicated by arrows.

**A**

```
                     C A R S P S G S G [N W S] Y S A G I D L W
            TG0081:  TGCGCGAGAAGTCCTAGTGGTAGTGGTAATTGGAGTTATTCTGCTGGAATTGATCTGTGG
Germline sequence:   TGCGCCAGAGCTACTGGTAGTCGTGGT                    TGCTGG ATTGATCTGTGG
                            IGHV1-23                                IGHD1    IGHJ1
```

**B**

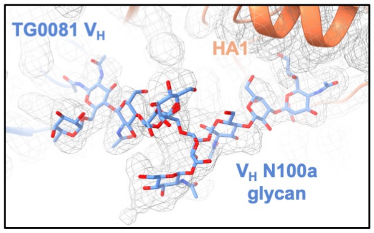


**C**

```
            45              61
IGHV1-1     LEYVASINSSGSSTYYAP
IGHV1-23    FEFVAQINPGGSYTGYAS
TG0081      FEFVAQINPGGSYTGYAS
```

**Figure S12. Sequence and structural analysis of TG0081. (A)** Amino acid and nucleotide sequences of the heavy chain V-D-J junction are shown for TG0081. Putative germline sequences and segments for *IGHV*, *IGHD*, and *IGHJ*, are indicated and colored in blue, purple, and red, respectively. Somatically mutated nucleotides are underlined. Intervening spaces at the V-D and D-J junctions are N-nucleotide additions. **(B)** Cryo-EM density maps (grey mesh) of H3N8 HA in complex with TG0081. **(C)** Sequence alignment of residues 45 to 61 of *IGHV1-1* (functional gene), *IGHV1-23* (pseudogene), and TG0081. Red: differences from *IGHV1-1*.
